# *Drosophila* nicotinic acetylcholine receptor subunits and their native interactions with insecticidal peptide toxins

**DOI:** 10.1101/2021.08.13.456240

**Authors:** Dagmara Korona, Benedict Dirnberger, Carlo N G Giachello, Rayner M L Queiroz, David-Paul Minde, Michael J Deery, Glynnis Johnson, Karin H Müller, Lucy C Firth, Fergus G Earley, Steven Russell, Kathryn S Lilley

## Abstract

*Drosophila* nicotinic acetylcholine receptors (nAChRs) are ligand-gated ion channels that represent a target for insecticides. Peptide neurotoxins are known to block nAChRs by binding to their target subunits, however, a better understanding of receptor subunit composition is needed for effective design of insecticides. To facilitate the analysis of nAChRs we used a CRISPR/Cas9 strategy to generate null alleles for all ten *nAChR* subunit genes in a common genetic background. We studied interactions of nAChR subunits with peptide neurotoxins by larval injections and styrene maleic acid lipid particles (SMALPs) pull-down assays. For the null alleles we determined the effects of α-Bungarotoxin (α-Btx) and ω-Hexatoxin-Hv1a (Hv1a) administration, identifying potential receptor subunits implicated in the binding of these toxins. We employed pull-down assays to confirm α-Btx interactions with the D*α*5, Dα6, D*α*7 subunits. Finally, we report the localization of fluorescent tagged endogenous Dα6 during nervous system development. Taken together this study elucidates native *Drosophila* nAChR subunit interactions with insecticidal peptide toxins and provides a resource for the *in vivo* analysis of insect nAChRs.

## Introduction

Global climate change and other factors are placing increasing demands on available agricultural land to deliver efficient, reliable and sustainable food production. Insecticides are important tools in securing yields of all major crops but need to be continually replaced to overcome resistance in target species and reduce environmental impacts. In addition, new insecticides must have low toxicity to non-target species, particularly the major pollinators essential for agriculture. A large class of insecticide targets are neurotransmitter receptors such as the nicotinic acetylcholine receptors (nAChRs) located in synaptic plasma membranes (Ihara et al., 2020). These pentameric cys-loop ligand-gated ion channels consist of either only α- subunits or α- and β-subunits, with ligand binding sites located between two α-subunits or between α- and β-subunits. Most insect genomes, including that of the highly tractable *Drosophila melanogaster* model, harbour ten highly conserved subunit genes that assemble in various combinations to form the active receptors.

An essential pre-requisite for effective design of new insecticides targeting these receptors is understanding the subunit composition of nAChRs and their distinctive binding properties. For many reasons, including low expression in endogenous tissues or difficulties in expressing insect receptors in heterologous systems, the characterisation of functional insect receptors has been challenging (Perry et al., 2021; Zuo et al., 2021; Salgado, 2021). Even in the tractable *D. melanogaster* insect model, there has been no systematic isolation of mutations in *nAChR* subunit genes, until recently, when Perry and colleagues described the generation of a new set of null mutations in nine out of the ten *D. melanogaster* subunit genes (Perry et al., 2021). These mutations, however, were generated in different genetic backgrounds necessitating additional work to assay background sensitive phenotypes such as neural or behavioural defects.

Several classes of insecticide, the most effective being those in the neonicotinoid and spinosad class, have been shown to bind insect nAChRs highly selectively to block their functions (Chambers et al., 2019; Houchat et al., 2019). Recently, the binding affinity and the positive allosteric effects of ω-Hexatoxin-Hv1a (Hv1a) peptide on nAChRs has been demonstrated (Chambers et al., 2019) and this spider venom peptide is well known for its insecticidal effects. In addition, other peptide toxins, such the snake venom constituent, α-Bungarotoxin (α-Btx), have been widely used to probe nAChR functions, however whether α-Btx harbours a selective insecticidal property is currently unknown. Alpha-Btx is a 74 amino acid peptide that binds irreversibly to nAChR α-subunits in different species, including *D. melanogaster,* although the exact subunit composition of target receptors is not fully understood (Schmidt-Nielsen et al., 1977; Dellisanti et al., 2007; Dacosta et al., 2015). Landsdell and co-workers have shown binding of α-Btx to *D. melanogaster* Dα5, Dα6, and Dα7 subunits in a heterologous S2 cell expression system (Lansdell & Millar, 2004; Lansdell et al., 2012) and the amino acid sequence of these subunits show strong similarity across their ligand*-*binding domains (LBD).

The lipid bilayer surrounding nAChRs is known to be essential for structural integrity, stability and ligand binding (Dacosta et al., 2013). However, this lipid requirement can make analysis of membrane protein complexes challenging. The development of methods for extracting membrane proteins from lipid bilayers using detergents and introducing them into artificial lipid nanodiscs has facilitated a much better characterisation of receptor-ligand interactions (Denisov & Sligar, 2016). The use of detergents generally used to solubilize membrane proteins, however, leads to destabilisation, aggregation and misfolding and are therefore not compatible with this type of analysis (Loo et al., 1996). Styrene maleic acid lipid particles (SMALPs) allow detergent-free extraction of membrane proteins in their local lipid environment and provide a promising technique for investigating receptor-ligand interactions under native conditions (Lee et al., 2016). This is particularly important since loss of lipids surrounding membrane proteins can lead to changes in measured binding affinities (Martens et al., 2018; Gault et al., 2020). The combination of detergent free SMALPs extraction coupled with mass spectrometry analysis provides a potential route for characterising native membrane receptor complexes (Sobotzki et al., 2018; Kalxdorf et al., 2021).

Here we report the results from a combined genetic and biochemical analysis of *D. melanogaster* nAChRs *in vivo*. Using CRISPR/Cas9 genome engineering we generated new null mutations for all ten receptor subunit genes in a uniform genetic background as well as introducing a fluorescent protein tag into the *nAChRα6* locus. We show that the null mutants in all seven α-subunit genes and two of the three β-subunit genes are viable and fertile, although we find mild morphological defects and some neurological impairment. Mutation of the remaining subunit gene, *nAChRβ1,* is recessive lethal. All nine of the viable null mutants were used to demonstrate a novel selective insecticidal effect of α-Btx on the *nAChRα5, nAChRα6* and *nAChRα7* subunits. We also applied the insecticidal Hv1a peptide to the viable null mutants, showing resistance with two subunit gene mutants: *nAChRα4* and *nAChRβ2.* In our biochemical studies we analysed receptor-ligand interactions in native conditions using SMALPs to verify the *in vivo* receptor subunit composition of the α-Btx binding target in adult neural tissue from wild-type and receptor subunit mutants. Our analysis revealed binding of α- Btx to receptors containing Dα5, Dα6 and Dα7 subunits with the analysis of mutants in these subunits genes indicating heterogeneity in α-Btx binding nAChRs. Furthermore, we have identified specific glycosylation sites in Dα5 and Dα7 subunits which are known from other studies to play a critical role in α-Btx binding affinity (Dellisanti et al., 2007; Rahman et al., 2020). Localization studies with the Dα6 subunit tagged at the endogenous locus with a fluorescent reporter shows expression at different developmental stages in specific neuronal cells, including the Kenyon cells of the mushroom bodies, a known site of α-Btx-binding.

## Results

### New *D. melanogaster* nicotinic acetylcholine receptor subunit gene mutations

To investigate the role of individual nAChR subunits we used CRISPR/Cas9 to generate deletion mutations in each of the seven α-subunit and three β-subunit genes. All of the mutations were generated in virtually identical genetic backgrounds using nanos-Cas9 sources on the second or third chromosome of otherwise genetically homogeneous fly lines. In brief, for each gene we targeted exons shared between all predicted isoforms, close to the N-terminus of the protein. In order to disrupt each coding sequence and facilitate screening we introduced a visible fluorescent marker, DsRED under control of the eye-specific 3xP3 promoter at the targeted locus. Positive lines were confirmed by PCR and sequencing, and subsequently the DsRED marker was excised from the genome by Cre-Lox recombination.

For nine out of ten subunit genes we established homozygous viable and fertile stocks, the exception was the *nAChRβ1* gene which proved to be recessive lethal. Although all the other lines are viable, we noticed that most of the mutants, but particularly *nAChRα1, nAChRα2, nAChRα5* and *nAChRβ3*, exhibited a curled abdomen phenotype that is most prominent in males (approximately 25, 20, 15 and 15 % respectively, Figure 1A). It is possible that this phenotype is a result of defects in neural control of abdominal muscles and it is interesting to note that a previous analysis of an *nAChRα1* allele reports reduced male courtship and mating (Somers et al., 2017). Since nAChRs are mostly found in the nervous system, we carried out basic climbing assays on the null alleles to assess potential locomotor defects (Figure 1B, Appendix-table 1). We saw little or no impact on the locomotor activity of ten day old flies with *nAChRα4, nAChRα5, nAChRα7, nAChRβ2* or *nAChRβ3* homozygous mutants, however, deletions of *nAChRα1*, *nAChRα2* and *nAChRα6* showed 50-60 % reductions in climbing ability compared to wild-type. In addition, the *nAChRα3* null mutant and heterozygotes for *nAChRβ1* exhibited a severe reduction in locomotor activity to less than 40 % of wild-type (22% and 34% respectively).

**Figure 1.**
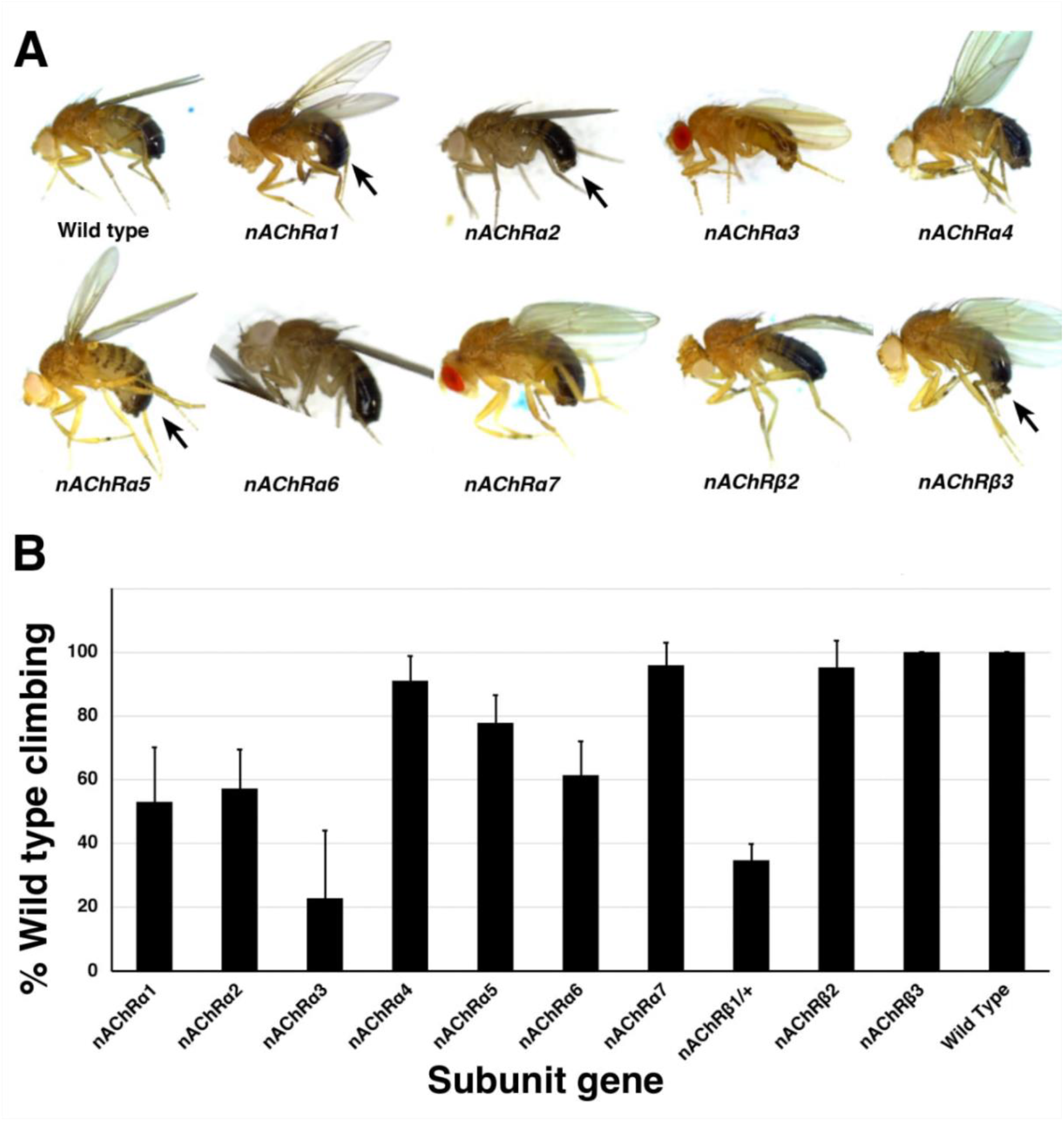
Morphological and locomotor phenotypes in *nAChR* subunit mutants. **(A)** Adult males from indicated *nAChR* subunit null mutants. Arrows indicate strong curled abdomen phenotypes. (**B)** Graph of locomotor activity determined in climbing assays as a percentage of wild type. Error bars represent standard deviation from 5 replicates.

Taken together, we report the generation and validation of null mutations in all ten *D. melanogaster nAChR* subunit genes, with mild morphological defects associated with most of the new alleles and impaired locomotion observed with some mutants.

### Distinct nAChR subunits mediate interactions with ω-Hexatoxin-Hv1a and α- Bungarotoxin

In order to investigate the selective contribution of each *nAChR* subunits to toxin binding *in vivo*, we injected 3^rd^ instar larvae from the homozygous *nAChR* null mutants with either ω- Hexatoxin-Hv1a (Hv1a) or α-Bungarotoxin (α-Btx) dissolved in PBS. As a control, injections of PBS alone (vehicle) were performed in parallel, and all larvae survived the injection procedure and showed no detectable defects. Larval injection of 2.5 nmol/g Hv1a induced locomotor paralysis and full lethality in the control groups (*w^1118^*, *THattP40* and *THattP2*, Figure 2A, Appendix-table 2). Survival was quantified as the percentage of pupae formed after injection. Hv1a did not result in full lethality with *nAChRα4* and *nAChRβ2* homozygous mutants, since both showed an increase in survival to 42±22% (One-way ANOVA followed by Bonferroni’s test, *P=*0.0035, Figure 2A). Mortality in all the other null mutants was comparable to controls (*P>*0.9).

**Figure 2.**
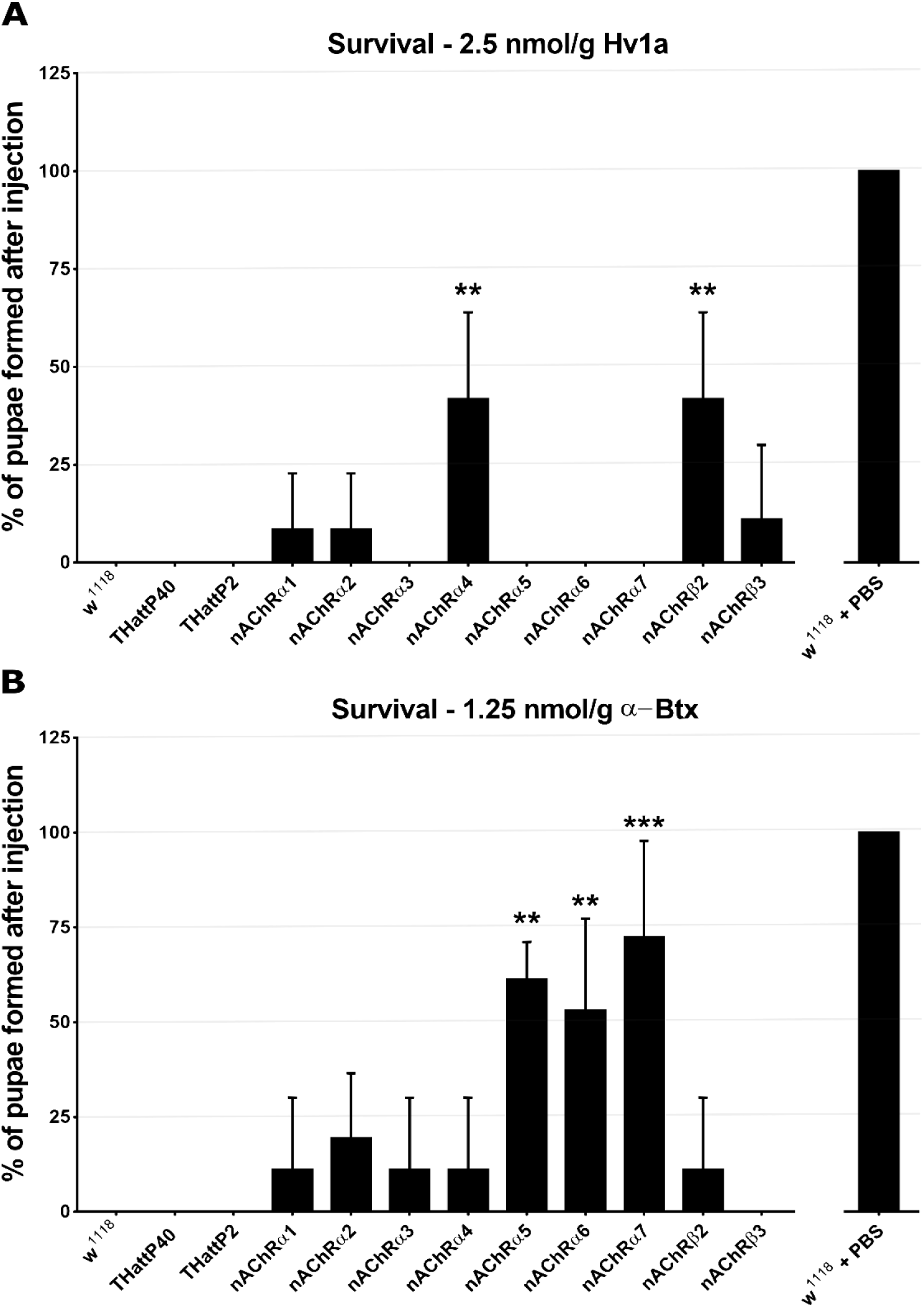
ω-Hexatoxin-Hv1a and α-Bungarotoxin target different nAChR subunits. **(A)** Bar graph of the survival rate, measured as the percentage of pupae formed, following larval injection of 2.5 nmol/g Hv1a in the indicated homozygous lines. Mean ± SD of 3 independent replicates of 10 larvae per replicate. *\*\*P=*0.0035 (one-way ANOVA (F_(11,24)_=4.99, *P*=0.0005 with Bonferroni’s multiple comparisons test). 3 independent replicates in each group (10 injected larvae in total). **(B)** Survival rate following larval injection of 1.25 nmol/g α-Btx. Mean ± SD of 3 independent replicates of 10 larvae per replicate. \**\*P<*0.001, \*\*\**P*=0.0001 (one-way ANOVA (F_(11,24)_= 7.921, *P*<0.0001, followed by Bonferroni’s multiple comparisons test). 3 independent replicates in each group (10 injected larvae in total). *w^1118^* is the wild type base stock, *THattp40* and *THattP2* are the Cas9 lines used to establish the mutants, *w^1118^* + PBS represents the injection control.

We also observed significant toxicity following injection of 1.25 nmol/g α-Btx, with larvae exhibiting a progressive reduction in locomotion until stationary, resulting in developmental arrest and death. We found that α-Btx induced lethality is drastically reduced in the *nAChRα5*, *nAChRα6* and *nAChRα7* subunit mutants, with the survival rate significantly increased from 0% (controls) to 61±10% (*P=*0.001), 53±24% (*P=*0.0051) and 72±25% (*P=*0.0001) respectively (One-way ANOVA followed by Bonferroni’s test, Figure 2B).

Together, these results indicate that Hv1a and α-Btx do not share the same binding target and differentially interact with the nAChR subunits *in vivo*. Since α-Btx showed a novel insecticidal effect on nAChRs we further examined its interactions biochemically.

### Forming SMA-lipid particles (SMALPs) of ring-like nAChR complex structures

In order to take advantage of our new receptor subunit mutants for the biochemical analysis of native nAChR functions, we examined the composition of the receptors responsible for binding α-Btx. To address the functionality of *D. melanogaster* nAChRs isolated from endogenous membranes, we utilised detergent-free SMALPs extraction to characterise the interaction between receptor native lipid discs and the α-Btx toxin (Figure 3A).

**Figure 3.**
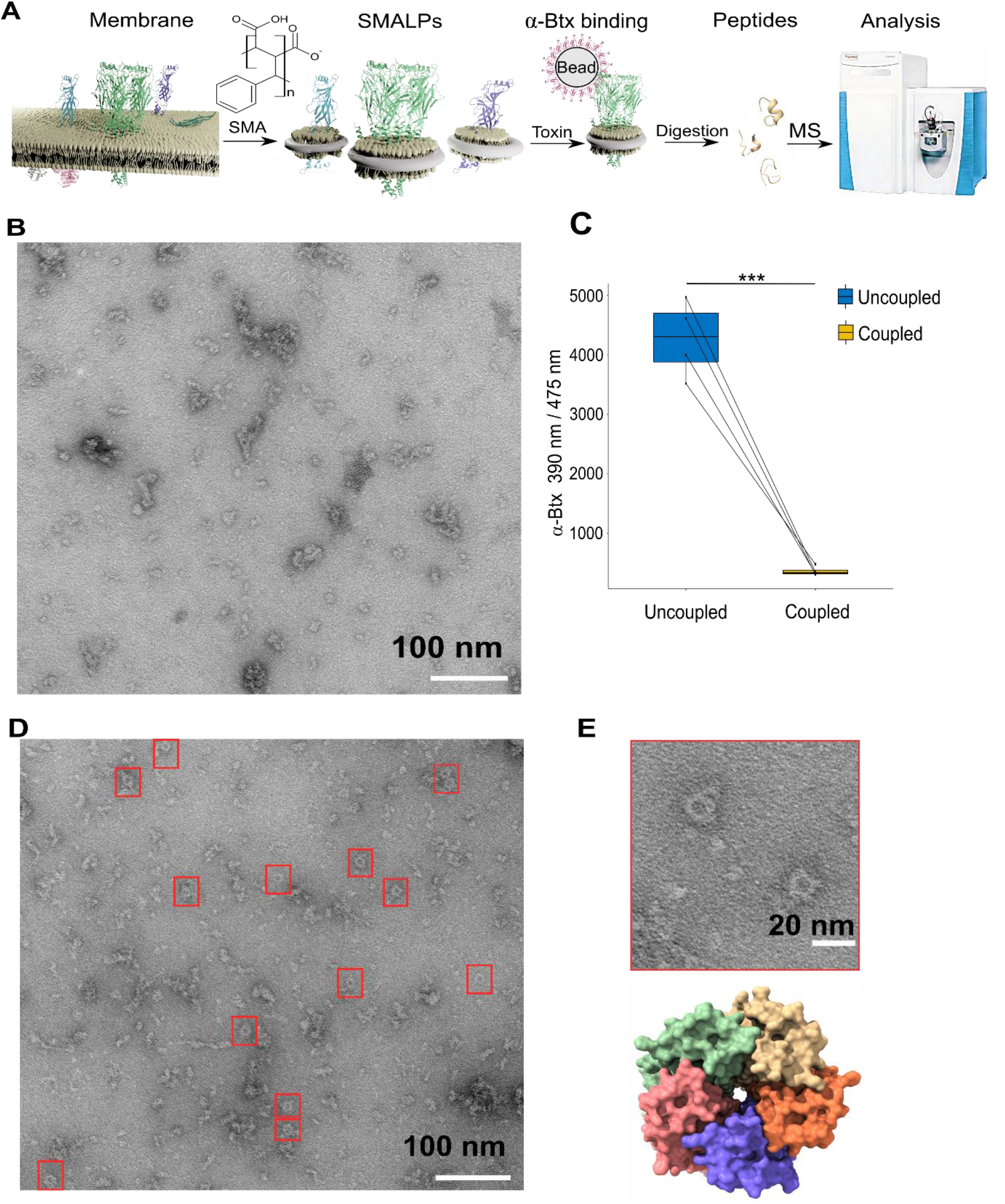
Forming styrene maleic acid lipid particles (SMALPs). **(A)** Schematic representation of the SMALPs extraction and nAChRs pull-down for mass spectrometric analysis. (**B)** Negative staining of extracted SMALPs by transmission electron microscopy. Scale bar 100 nm. (**C)** Fluorescence signal of uncoupled α-Btx in solution before and after coupling to affinity beads (two-tailed t-test, \*\*\**P*<0.001, n=4). (**D, E)** Negative staining of extracted SMALPs after α-Btx pull-downs. Ring-like protein structures are boxed (Scale bar = 100 nm) with an example in the magnified image (Scale bar = 20 nm). A top view of the nAChR structure from PDB entry 4HQP is shown for reference.

In brief, we prepared membrane extracts from adult *D. melanogaster* heads (Depner et al., 2014) and generated lipid particle discs by solubilising the membrane extracts with the SMA copolymer. We used affinity beads coupled to α-Btx (Wang et al., 2003; Mulcahy et al., 2018) to enrich for nAChRs in the SMALP preparations that bound to the toxin, and performed mass spectrometric analysis of tryptic peptides generated from the enriched preparations. In parallel we processed membrane extracts without SMALP and with SMALP extracts enriched with beads alone.

We first determined whether membrane protein discs are formed from enriched membranes using the SMA copolymer. We prepared membrane enriched fractions from adult heads, solubilized these with SMA and separated the insoluble particles from the lipid discs by ultracentrifugation. We negatively stained the SMALP preparations and imaged them with transmission electron microscopy (TEM), observing irregular discs of varying shapes and sizes, with clusters containing different numbers of discs (Figure 3B).

Membrane receptors often have a unique shape in TEM images and the five subunits of a nAChRs is expected to form a ring-like structure, suggesting that the receptors are extracted as a complex. However, we did not observe pentameric ring-like structures perhaps suggesting that nAChRs are of low abundance and that analysis may benefit from enrichment. We coupled α-Btx to affinity beads to enrich nAChR complexes that bind the toxin in SMALP preparations (Figure 3C). In contrast to the unenriched samples, TEM images of the enriched preparations showed increased numbers of ring-like structures of 15 nm in diameter (Figure 3D, E). Thus our TEM analysis shows an increased number of ring-like membrane complexes in the SMALP preparations which are likely to be nAChRs.

### Efficient SMALPs extraction allow to study nAChR subunits solubility

To assess to what extent the SMA copolymer solubilized nAChRs, we performed a bottom-up proteomics analysis to identify receptor subunits. Membrane preparations were solubilized in buffer with or without SMA, and affinity beads with or without α-Btx were used to assess ligand-binding to nAChR subunits. Comparing the number of proteins identified in samples solubilized either with or without 5% SMA, we observed a significantly increased identification rate of proteins dissolved in SMA by equal numbers of MS/MS spectral counts (two-tailed t-test*, P*<0.01, Figure 4A and non-significant, Figure 4B). This indicates that mass spectrometer performance was comparable during the measurements.

**Figure 4.**
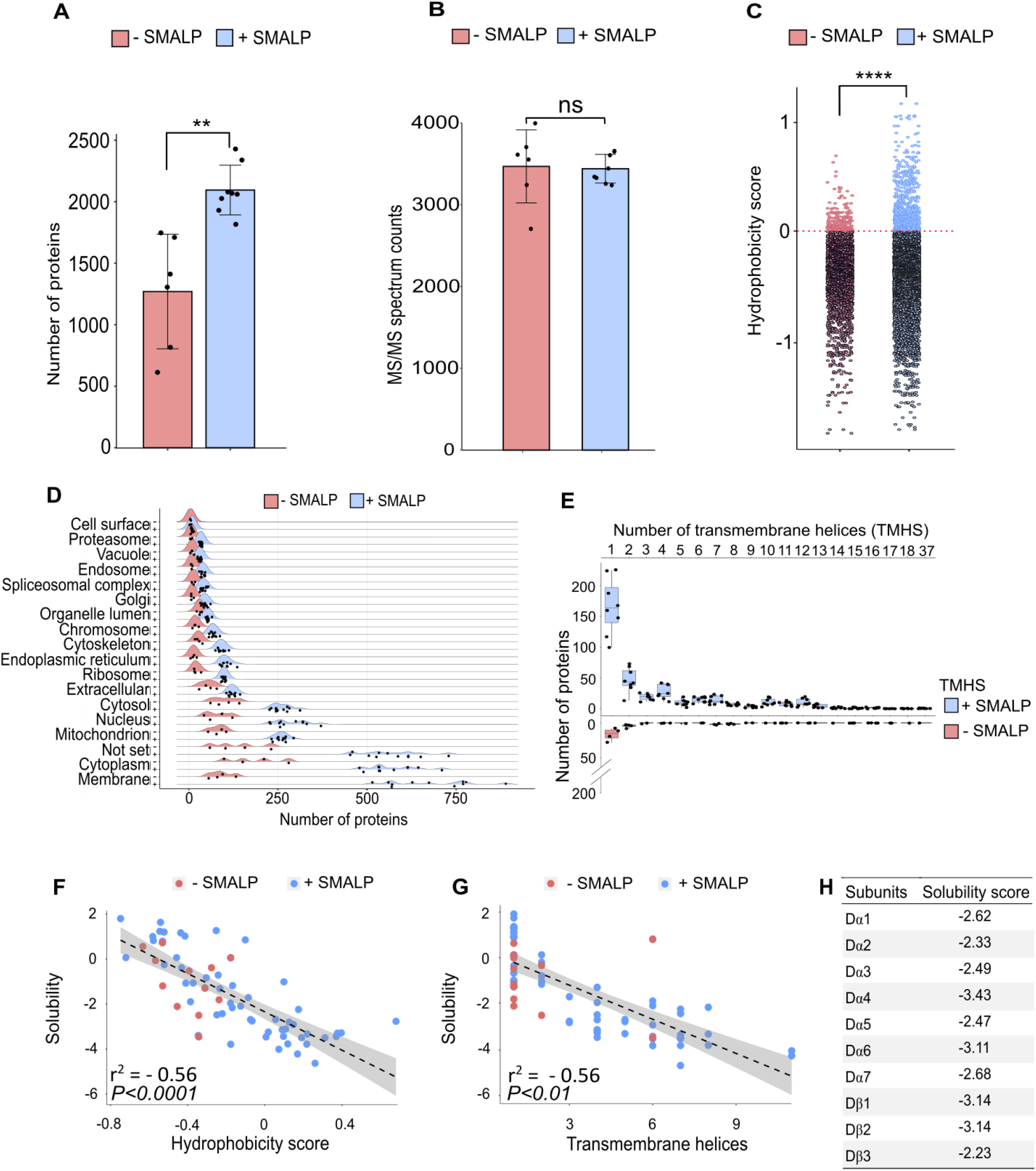
Identification of proteins enriched by SMALP extraction. **(A)** Number of identified proteins in affinity pull-down samples solubilized with or without SMA, two-tailed t-test, \*\**P*<0.01, n=6 or 8 replicates per condition. (**B)** MS/MS spectrum counts from samples solubilized with or without SMA, ns = not significant after two-tailed t-test with n=6 or 8. (**C)** Calculated hydrophobicity score of amino acid residues found in protein sequences obtained with and without SMA solubilisation, \*\*\*\**P*<0.0001, two-tailed t-test, n=3 per condition. (**D)** GO term (cellular compartment) enrichment of proteins identified with and without SMA solubilisation, n=4 or 11. (**E)** Predicted numbers of proteins containing transmembrane helices obtained with or without SMA solubilisation, n=4 or 8. (**F, G)** Analysis of solubility and hydrophobicity of receptors identified with and without SMA solubilisation (r^2^= -0.56, *P*<0.0001, n=4) and of transmembrane receptor helices (r^2^=0.56, *P*<0.01, n=4 ). (**H)** Solubility score of individual nAChR subunits.

Sequences of membrane spanning segments of nAChR subunits, which are in close contact to the hydrophobic lipid environment, are largely composed of nonpolar side chains. Determining the average of hydrophobicity of identified protein sequences revealed significantly increased numbers of proteins with a positive hydrophobicity score in samples solubilized in SMA (two-tailed t-test*, P*<0.0001, Figure 4C), indicative of enrichment of membrane proteins. An analysis of Gene Ontology (GO) slim terms supports the conclusion that the SMALP preparations are enriched of membrane embedded and associated proteins (Figure 4D), and that these are not limited to plasma membrane proteins. In the SMA-enriched samples we found enrichment for proteins annotated with metabolic and catalytic activity terms and also enhanced response to biological stimuli (Figure 4-figure supplement 1A, B), highlighting the recovery of membrane-associated proteins.

Next, we focused on identified membrane proteins predicted to contain transmembrane helical (TMH) domains and found an increased number of proteins containing TMHs in SMA solubilized samples (Figure 4E). While the majority of these proteins contained a single TMH domain, we identified Piezo, a mechanosensory ion channel protein containing 37 predicted transmembrane helices. Both α- and β-nAChR subunits contain four TMH domains and could be solubilized in SMA. The number of β-barrel membrane spanning proteins identified was also significantly increased by SMA extraction (two-tailed t-test, *P*<0.0001, Figure 4-figure supplement 1C). In addition, palmitoylated lipid anchor modifications to nAChR subunits has been shown to be important for receptor assembly into membranes and the formation of functional complexes (Alexander et al., 2010).

Comparing samples solubilized with and without SMA showed a significantly increased identification of proteins which are predicted to be palmitoylated and myristoylated (two-tailed t-test*, P*<0.0001, Figure 4-figure supplement 1D, E). In contrast, membrane proteins that are predicted to contain a glycosylphosphatidylinositol (GPI)-anchor are equally solubilized in both conditions (two-tailed t-test, non-significant, Figure 4-figure supplement 1F).

Focusing on the membrane receptors solubilized by SMA, we analysed the amino acid sequence properties of identified proteins and calculated an overall solubility score (Sormanni et al., 2015; Sormanni et al., 2017). Comparing the solubility to the hydrophobicity showed a calculated R^2^ of 0.56 (Figure 4F). Sequences with a score greater than 1 are highly soluble receptors and those less than minus -1 are difficult to solubilize. As a result, samples solubilized in SMA contain more receptors, which are difficult to solubilize. These receptors are more hydrophobic and contain larger numbers of TMH domains (Figure 4G). Calculating an average solubility score of -2.76 for nAChR sequences indicates that difficult to solubilize subunits are successfully recovered with SMA (Figure 4H).

Taken together, these analyses confirm that SMA solubilizes nAChR complexes in a state suitable for subunit identification by mass spectrometry and suggests that α-Btx interactions can be studied with SMALP preparations.

### Three nAChR α-subunits are targets of α-Btx

To explore native nAChR subunit interactions with α-Btx we searched for peptides from subunit ligand-binding and cytoplasmic domains, identifying the Dα5, Dα6 and Dα7 subunits in the α-Btx affinity bead pull-downs (Figure 5A and B, Appendix-table 3). Several other nAChR subunit peptides could be identified in the negative controls performed without coupling α-Btx to affinity beads (Appendix-table 4). The sequences of the ligand-binding domains of the Dα5, Dα6 and Dα7 subunits are very similar (avg. 95.49 %) and we identified peptides common to all three subunits (Figure 5-figure supplement 2A) as well as unique peptides within their cytoplasmic domains (Figure 5-figure supplement 2B). However, we found no evidence of peptides mapping to TMH domains. The ligand-binding domain of α- subunits show structural similarity across different species (Figure 5-figure supplement 3A) and by mapping the identified peptides to known structures we concluded they are most likely outside of the α-Btx binding sites (Figure 5-figure supplement 3B).

**Figure 5.**
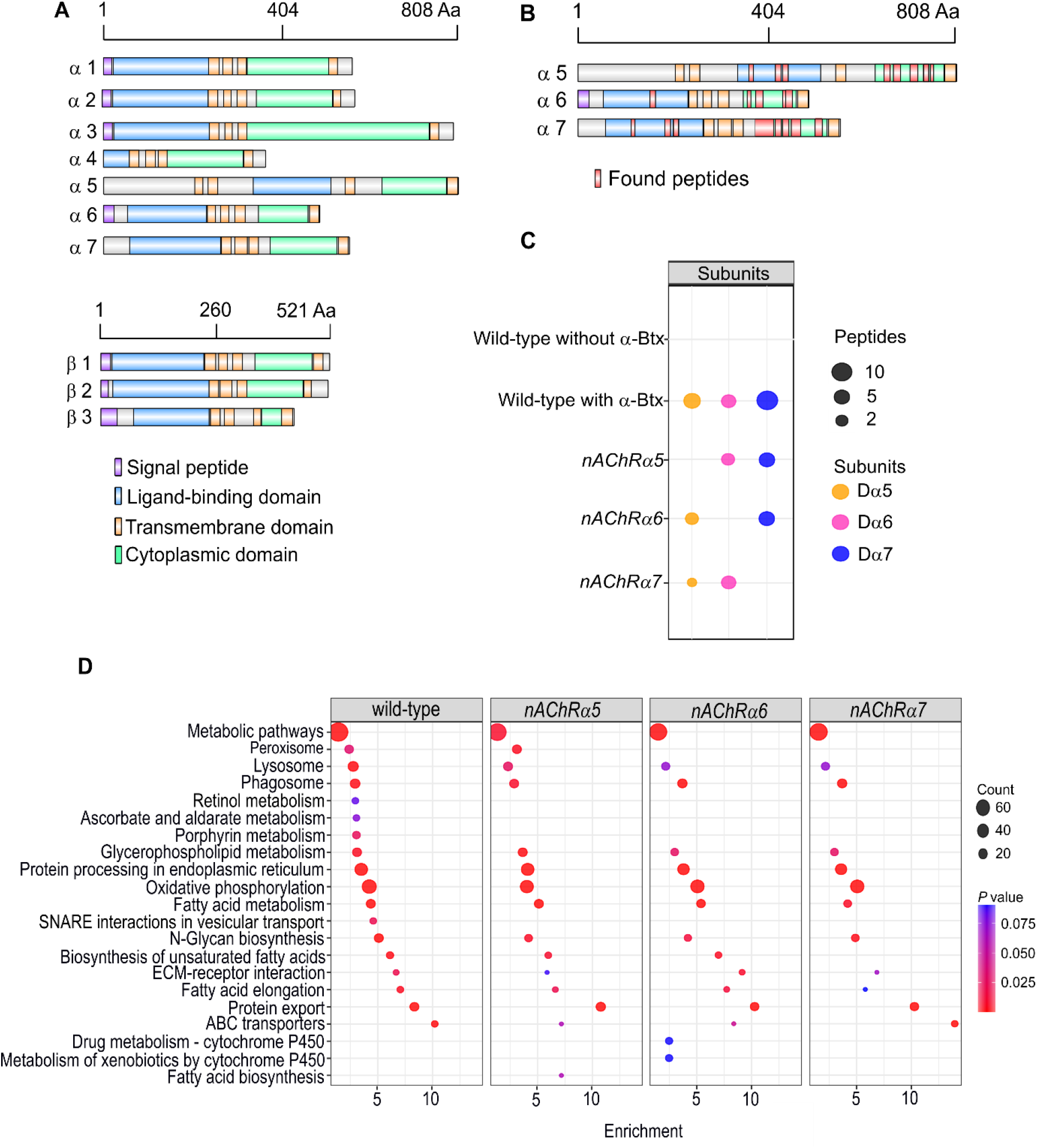
Three nAChR α-subunits are binding to α-Bungarotoxin (α-Btx). **(A)** Graphical representation of ten nAChR subunits. The position of protein domains and signal peptides are shown. (**B)** Identified peptides of Dα5, Dα6 and Dα7 nAChR subunits in pull-downs using α-Btx affinity beads. Found peptides in ligand-binding and cytoplasmic domain are highlighted in red. (**C)** Numbers of identified unique peptides in wild-type pull-downs using affinity beads in absence and presence of α-Btx, n=3. Deleting *nAChRα5, nAChRα6, nAChRα7* and performing pull-downs identified unique peptides of nAChR subunits suggesting that functional complexes can be formed in null alleles, n=3. (**D)** KEGG pathway enrichment analysis of pull-downs in wild-type and *nAChRα5, nAChRα6, nAChRα7* null alleles, Fisher’s exact test, n=3. Protein counts with *P* values of enriched pathways are shown. *P* values of ≤ 0.05 are to be considered as strongly enriched with default threshold of 0.1.

To further characterize the role of the three α-subunits identified in α-Btx binding we generated SMALP preparations and performed α-Btx affinity bead enrichments with adult head preparations from homozygous null mutations for each of the *nAChRα5*, *nAChRα6* and *nAChRα7* subunit genes. With all three deletion mutants we observed, as expected, no detectable peptides from the missing subunit but could still identify peptides from the other two subunits (Figure 5C).

We compared the repertoire of proteins identified with α-Btx enrichment in wild-type with those found in each of the three mutant lines to identify any changes in the representation of biological pathways annotated in KEGG (Kanehisa et al., 2020, Figure 5D). While the enrichments in wild-type and the mutants were broadly similar, we noticed a loss of proteins associated with cofactor/vitamin metabolism, particularly retinol and ascorbate, in all three of the mutants as well as proteins associated with vesicular transport. It is possible that these pathway changes represent alterations in neurotransmitter production or trafficking. Interestingly, we also noticed specific enrichment of cytochrome P450 related pathways in the *nAChRα6* mutants, suggesting perturbation of neurotransmitter pathways.

In summary, our analysis indicates that a functional α-Btx binding nAChR involves the Dα5, Dα6 and Dα7 subunits. This is entirely in line with our genetic findings described above, where loss of each of these subunit genes conferred substantial resistance to α-Btx induced lethality.

### Glycosylation sites of nAChR subunits by α-Btx binding

We next examined glycosylation sites on nAChR subunits since these are known to have an important role in α-Btx binding affinity in other systems. For example, deglycosylation reduces α-Btx binding in human nAChRs by more than two orders of magnitude (Dellisanti et al., 2007) and α-Btx binding to loop C in *Torpedo californica* α-subunits is enhanced by N-glycosylation of sites in these regions (Rahman et al., 2020). To identify specific glycosylation sites in *D. melanogaster* nAChRs we first purified SMALP solubilized receptors with α-Btx affinity beads, digested them into peptides and enriched for glycopeptides using HILIC resin (Hägglund et al., 2004, Figure 6A). Site-specific identification of glycans on peptides by mass spectrometry is challenging (Fang et al., 2020) and often requires an additional deglycosylation step for glycopeptide measurement.

**Figure 6.**
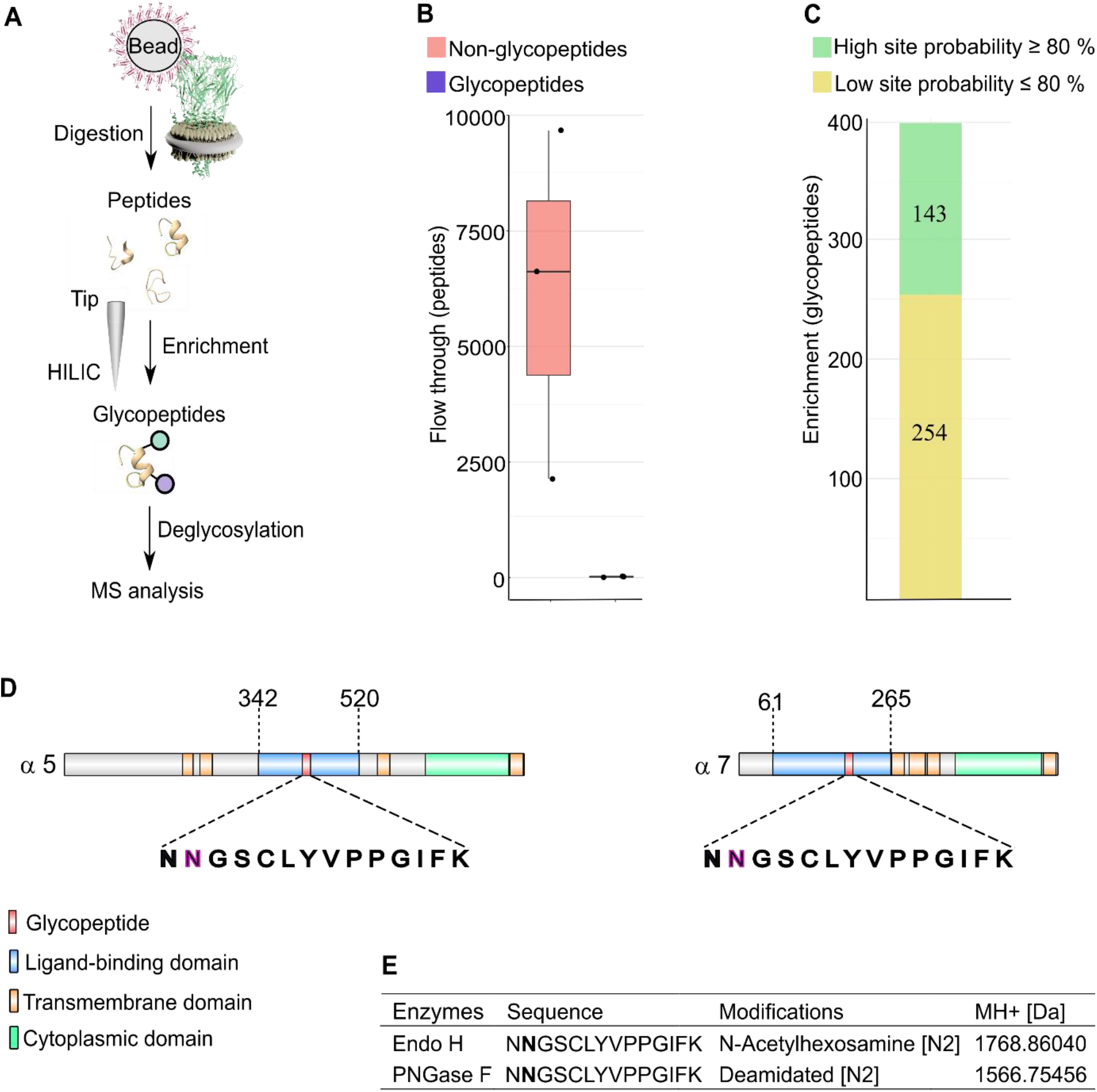
N-glycosylation sites in nAChR subunits. **(A)** Diagrammatic representation of nAChR subunit glycopeptide enrichment. Pull-downs with α-Btx affinity beads enrich for nAChRs and after tryptic digestion glycopeptides were enriched. Glycopeptides were deglycosylated with Endo H or PNGase F and analyzed my mass spectrometry. **(B)** Low numbers of glycopeptides (average 20) are detected in flow through fractions. (**C)** Numbers of identified glycopeptides according to site probabilities are shown (n=3). (**D)** Shared glycopeptide identified in the ligand-binding domain of Dα5 and Dα7, an N-linked glycosylated asparagine (N) residue is highlighted. (**E)** Deglycosylated peptide with either Endo H or PNGase F and contains either an N-acetylhexosamine or is deamidated on asparagine (N2). The two different modifications on the same peptide lead to a different monoisotopic mass (MH+ [Da]). Peptide contains an additional carbamidomethyl on cysteine (C5)

Deglycosylation of enriched peptides was carried out using two separate enzymes: Endoglycosidase H (Endo H), which cleaves asparagine-linked oligosaccharides to generate a truncated sugar molecule with one N-acetylhexosamine (HexNAc) residue, and the endoglycosidase PNGase F, which releases the entire glycan from asparagine residues and deaminates the sugar free asparagine to aspartic acid. While very few glycopeptides were observed in the flow through (an average 20 glycopeptides Figure 6B), we identified a total of 397 glycopeptides after enrichment and deglycosylation with Endo H or PNGase F (Figure 6C).

Shared glycopeptides from Dα5 and Dα7 nAChR subunits were identified after enrichment and deglycosylation with Endo H or PNGase F (Figure 6D). Deglycosylation with Endo H identified modified asparagine (N2) residues on the peptide (NNGSCLYVPPGIFK), which is predicted to be part of the Dα5 and Dα7 ligand-binding domains involved in α-Btx binding. This asparagine residue was modified with an N-acetylhexosamine (HexNAc) truncated sugar chain. Releasing N-glycans after deglycosylation by PNGase F enabled us to identify a deaminated asparagine residue in the same peptide. The monoisotopic mass of this peptide changed due to the different modifications on the asparagine residue (Figure 6E).

The genome of *Caenorhabditis elegans* encodes for at least 29 nAChR subunits (Jones et al., 2007). The alpha-type unc-63 subunit contains an N-linked HexNAc modified asparagine residue on position 136 (Kaji et al., 2007). Performing a multiple sequence alignment showed that this asparagine residue is conserved between insects and nematodes (Figure 6-figure supplement 4A). Comparing identified glycosylation sites of Dα5 and Dα7 subunits to known N-linked glycosylation sites of α-subunits from *T. californica, Danio rerio, Mus musculus* or *Homo sapiens* indicates that this site is not conserved between vertebrates and invertebrates (Figure 6-figure supplement 4B).

We also identified glycosylation sites in the Dα3 (ATKATLNYTGR) and Dβ3 (VVLPENGTAR) subunits after Endo H treatment but not with PNGase F treatment, suggesting they harbour a single N-linked HexNAc modified asparagine residue (Figure 6- figure supplement 4C).

Taken together these findings suggest that the Dα5 and Dα7 subunits are modified at asparagine residues in the α-Btx ligand-binding domain with an N-linked sugar chain.

### Localization of Dα6 nAChRs subunit in the brain

In order to examine the endogenous localization of an α-Btx binding receptor subunit we used CRISPR/Cas9 genome engineering to introduce in frame C-terminal fluorescence and epitope tags into the endogenous *nAChRα6* locus (Figure 7).

**Figure 7.**
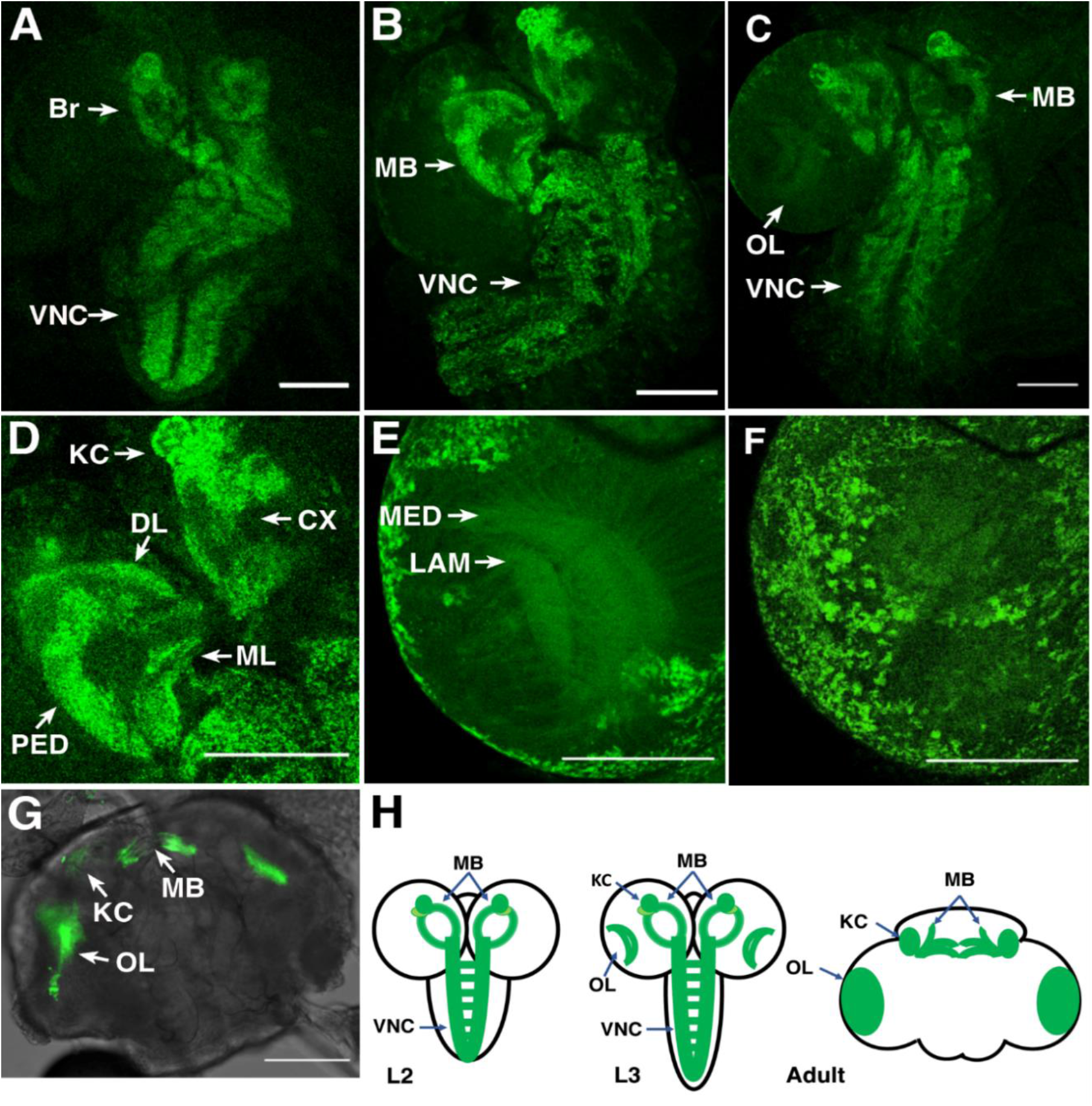
*In vivo* imaging of endogenously tagged Dα6 nAChR subunit. **(A-G)** Live imaging of fly brains carrying a C-terminal EGFP fusion into the endogenous *nAChRα6* locus. **(A-C)** Dα6 subunit in 2^nd^, early and late 3^rd^ instar larvae brain, respectively. Visible localization in ventral nerve cord (VNC), mushroom bodies (MB), and optic lobes (OL). Scale bar = 100 µm**. (D)** Dα6 subunit in mushroom bodies of 3^rd^ instar larvae with detectable fluorescence signal in Kenyon cells (KC), calyx (CX), peduncle (PED), dorsal lobes (DL) and medial lobes (ML). Scale bar = 100 µm**. (E)** Dα6 subunit was observed in developing optic lobes, lamina (LAM) and medulla (MED) of later 3^rd^ instar larvae. Scale bar = 100 µm. **(F)** Dα6 subunit on the external structures of developing lobes in later 3^rd^ instar larvae. Scale bar = 100 µm. **(G)** Dα6 subunit in adult fly brain, strong signal detected in mushroom bodies (MB) and optic lobe (OL). Scale bar = 100 µm**. (H)** Schematic summary of Dα6 subunit expression during different developmental stages, 2^nd^ and 3^rd^ instar larvae and adult fly, (L2, L3 and Adult, respectively) in which the green lines indicate the localization of the Dα6 subunit

The resulting line is homozygous viable and fertile, and shows no apparent phenotypes. We live imaged the unfixed brains of larvae and adults homozygous for the tagged line using confocal microscopy. In 2^nd^ instar larvae we observed low level well-distributed fluorescence signal throughout the ventral nerve cord (VNC), including on commissural axons, and in the developing brain (Figure 7A).

By early L3, we found more defined localization in the VNC and developing mushroom bodies (Figure 7B and D), particularly noticeable in the Kenyon cells, a known site of α-Btx binding (Su & O’Dowd, 2003). Localization in larval mushroom bodies continued to evolve, with defined expression in the Kenyon cells, calyx, peduncle, dorsal and medial lobes as well as the medulla and lamina of the emerging optic lobes (Figure 7C, E). We also observed localisation to a number of cell bodies overlying the optic lobes (Figure 7F).

Finally, in the adult brain, expression was largely restricted to the mushroom bodies particularly the Kenyon cells and connections across the midline between the β and γ lobes and the optic lobes (Figure 7G). The temporal localization of Dα6 subunit in the CNS is summarized in schematic form (Figure 7H).

## Discussion

Elucidation of complex insect nAChRs heterogeneity will lead to a better understanding of selective insecticidal effects. We present a new set of null mutations in all *D. melanogaster nAChR* subunit genes and investigated insecticidal peptide toxin effects on wild-type and receptor subunit mutant larvae. Utilising biochemical approaches with SMALP pull-downs we characterised toxin binding and subunit composition of native nAChR complexes.

Our genome engineering approach generated viable and fertile mutations in nine out of the ten subunit genes encoded in the *D. melanogaster* genome and is largely concordant with the recently described work by Perry and colleagues (Perry et al., 2021). In both studies, null mutations in the *nAChRβ1* gene were inviable as stocks. We add to the previous work by generating viable mutations in *nAChRα5*. We observed some minor morphological defects in some of the null mutants especially in *nAChRα1, nAChRα2, nAChRα5* and *nAChRβ3* as well as locomotor defects with some alleles, particularly severely in *nAChRα3* homozygotes and *nAChRβ1* heterozygotes. The locomotor defects we observed are in agreement with previously reported neuronal phenotypes with *nAChR* subunit genes, including sleep disruption, defective jump response, memory impairment or locomotor defects (Fayyazuddin et al., 2006; Rohde et al., 2016; Somers et al., 2017; Tackenberg et al., 2020).

We used the nAChR null mutants to study insecticidal effects of the Hv1a peptide on viability after injection into larvae and investigated whether α-Btx has any insecticidal properties. As described by Chambers and colleagues, we confirm that Hv1a effects nAChRs (Chambers et al., 2019) and our analysis shows that the Dα4 and Dβ2 subunits are involved in the insecticidal response. We show for the first time that α-Btx has selective insecticidal effects against the Dα5, Dα6 and Dα7 subunits, which we further characterized at the biochemical level.

The pharmacology of Hv1a and α-Btx binding has been shown to be distinctive (Chambers et al., 2019), correlating with our demonstration that these two peptide toxins mediate their effects through different receptor alpha subunits. Furthermore, resistance to neonicotinoid insecticides, which interact most strongly with Hv1a binding, has been associated with Dβ2 (Perry et al., 2008; Perry et al., 2021), consistent with the involvement of this subunit in the response to Hv1a. However, no resistance to neonicotinoids was seen in *D. melanogaster* carrying a *nAChRa4* gene deletion (Perry et al., 2021), which could be explained if neonicotinoids act at multiple receptor classes. Multiple binding sites for the neonicotinoid imidacloprid can be resolved in equilibrium binding assays in many insect species (Xu et al., 2010) and by binding kinetics in flies (Liu & Casida, 1993).

Resistance to spinosad is strongly associated with Dα6 (Perry et al., 2021), and spinosad binding is much more sensitive to the action of α-Btx than to the action of neonicotinoids (Chambers et al., 2019), again consistent with the involvement of this subunit with sensitivity to injected α-Btx and with the proposition that α-Btx and Hv1a act at distinct receptor classes.

nAChR subunits are known to be difficult to purify due to solubilisation issues (Cheng et al., 2015; Maldonado-Hernández et al., 2020) and the requirement for a lipid environment for ligand binding (Dacosta et al., 2013) makes it challenging to study these receptors in native conditions. We used the SMALPs extraction method for preparing membrane discs and enriched nAChRs via α-Btx affinity purification. Electron microscopy analysis indicated that receptor-like particles were recovered and these were substantially enriched by α-Btx pull-down. Mass spectrometry analysis showed an enrichment for the Dα5, Dα6 and Dα7 subunits in these preparations, which is concordant with our *in vivo* injection results and previous studies that characterised aspects of α-Btx binding (Lansdell & Millar, 2004; Wu et al., 2005; Lansdell et al., 2012).

To our knowledge this is the first report of the identification of a native endogenous α-Btx binding nAChRs. We note however, that we cannot determine from our analysis whether all three identified subunits are part of the same complex or if there are different receptors containing a subset of these subunits. Using chimeric receptors in a cell line system, Landsdell and colleagues reported that a combination of all three of these subunits show high affinity acetylcholine binding but α-Btx binding varied depending on receptor combinations, with Dα5 and Dα6 binding most strongly (Lansdell et al., 2012). In a prior study they implicated Dα6 and Dα7 (Lansdell & Millar, 2004). However, these assays were performed with 5HT3A- nAChR subunit fusions, here we provide strong evidence that these three subunits bind to α- Btx *in vitro* and *in vivo*.

In addition, glycopeptide enrichment showed site specific glycosylation modifications on the Dα5 and Dα7 nAChR subunit ligand binding domains. The unique lipid environment and glycosylation sites of nAChR α-subunits from the electric ray, *T. californica*, were found to be important for α-Btx binding activities (Quesada et al., 2016; Rahman et al., 2020), and structural studies support this conclusion (Dellisanti et al., 2007). Our work supports the view that there is a role for Dα5 and Dα7 glycosylation modifications in the recognition of α-Btx in *D. melanogaster*.

Our localization studies with fluorescence tagged endogenous Dα6 subunit showed relatively restricted expression in the brain and ventral nerve cord, with prominent expression in the Kenyon cells of the mushroom body, all known regions. The expression of Dα6 in Kenyon cells across development is in line with a proposed role for this subunit in memory plasticity, along with other α-subunits including Dα5, in mushroom body output neurons (Barnstedt et al., 2016). Thus, it is possible that retention of α-Btx binding in the absence of Dα6 may simply reflect its restricted localisation. In contrast, it is clear that Dα6 plays a major and specific role in binding to the insecticide spinosad in *D. melanogaster* since mutations in this subunit are highly resistant to the toxin (Perry et al., 2015).

Localization studies of Dα6 nAChRs subunit fusion protein by confocal microscopy are largely consistent with recent reports of *nAChRα6* expression derived from expression reporters (Kondo et al., 2020), though these studies appear to indicate wider adult brain expression than we observed, perhaps reflecting a degree of translational control or limitations in the sensitivity of our live imaging. In conclusion, we identified ligand-binding subunit sites for a *D. melanogaster* nAChR antagonist with newly insecticidal effects. Our findings contribute to a better understanding of the role of nAChR subunits which interacts with insecticidal peptide toxins.

## Materials and methods

### Drosophila methods

Embryos were injected using standard procedures into the *THattP40* (*y^1^ sc v^1^ sev^21^; P{y^+t7.7^v^+t1.8^ nos-Cas9.R}attP40*) or *THattP2* (*y^1^ sc v^1^ sev^21^; P{y^+t7.7^ v^+t1.8^ nos-Cas9.R}attP2*) lines expressing *nos*-Cas9 (Bloomington *Drosophila* Stock Centre). Donor DNA (500 ng/μL) in sterile H_2_O was injected together with of gRNA plasmids (100 ng/μL) as described previously (Korona et al., 2020). Individually selected surviving adults were crossed to *w^1118^* and the progeny screened for DsRED fluorescence localized mostly to the eyes of transgenic flies: positive flies were balanced and homozygous stocks established where possible. The correct localization of the insert was confirmed via PCR and sequencing. Transgenic flies were assessed for the phenotype using bright field microscope. For tagging of *nAChRα6*, the stocks were additionally subjected to Cre-recombination for marker removal and several independent lines were verified by PCR. Some of these lines were screened for YFP fluorescence using confocal microscopy. From the YFP positive balanced stocks, the viable and fertile homozygote was established. Injections were performed by the Department of Genetics Fly Facility (https://www.flyfacility.gen.cam.ac.uk). All fly stocks were maintained at 25°C on standard cornmeal medium. Larvae of 2^nd^ and 3^rd^ stage were collected, and their brains were dissected according to standard protocols. Brains were mounted in glycerol and live imaged.

### Cloning of gRNAs and generation of donor vectors Construction of *nAChR* subunits null alleles

In order to generate individual *nAChR* subunits gene deletions the open reading frame (ORF) was disrupted by introducing a visible marker harbouring DsRED marker under eye specific driver 3Px3 using CRISPR/Cas9 technology as previously described (Korona et al., 2020). The targeted exons are shared between different isoforms and adjacent to the N-terminus to ensure the protein translated was interrupted. The insertion sites were designed *in silico* and optimal gRNAs were chosen (Appendix-table 5) that were tested against the injection strain and cloned into pCDF3. Briefly, target specific sequences were synthesized and either 5′-phosphorylated annealed and ligated into the *Bbs*I sites of pCDF3 precut with *Bbs*I. Positive clones were confirmed by sequencing.

For generation of donor vectors, firstly, homology arms were amplified on genomic DNA (Appendix-table 6) that, secondly, were used as a template to amplify the homology arms (Appendix-table 7) of the donor vector for CRISPR/Cas9 homologous recombination (HDR). The inserts with visible marker were amplified using as a template previously generated constructs (Korona et al., 2020) with appropriate primers. These fragments were used for Gibson assembly using Gibson Assembly Master Mix (New England Biolabs). PCR products were produced with the Q5 High-Fidelity 2X Master Mix (New England Biolabs). All inserts were verified by sequencing.

### C-terminal tagging of Dα6 nAChRs subunit fusion protein

For tagging of Dα6 nAChRs subunit the C-terminal fusion with FSVS fluorescent protein harbouring StrepII and 3xFLAG epitope tags (3xFLAG-StrepII-Venus-StrepII) was generated for CRISPR/Cas9 mediated genome engineering (Korona et al., 2017; Korona et al., 2020). Firstly, gRNAs were designed (Appendix-table 5) and tested against the genomic DNA sequence of injection strains. The oligonucleotides were phosphorylated and ligated into *Bbs*I pre-cut pCDF3. The positive variants were confirmed by sequencing.

The donor vector to generate protein fusion with fluorescent protein harbouring epitope tags was cloned in 2 steps strategy by creating initially (A) nAChRα6-FSVS donor and then adding the removable marker to generate (B) nAChRα6-FSVS-loxP-3PX3_DsRED_loxP donor vector. At first, the homology arms were enriched on genomic DNA (Appendix-table 6) and used to amplify homology arms for donor vector nAChRα6-FSVS (Appendix-table 7) that was assembled using Gibson Assembly® as described above. The FSVS tag was amplified on previously generated constructs (Korona et al., 2017) with appropriate overlapping oligonucleotides ( Appendix-table 7). The construct was confirmed by Sanger sequencing and used as a template to generate donor vector with removable marker. The PCR fragments harbouring homology arms and FSVS tag were amplified on nAChRα6-FSVS construct, whereas the 3PX3-DsRed with adjacent loxP sites was amplified using earlier generated constructs (Korona et al., 2017). The final donor vector was generated using Gibson Assembly® as described above and positive variants were confirmed by sequencing.

### Confocal microscopy

Localization of FSVS-tagged (3xFLAG-StrepII-Venus-StrepII) Dα6 nAChRs subunit was visualised in dissected larvae brains via monitoring the YFP fluorescence (Venus). Briefly, the larval brains were dissected and mounted in glycerol for live imaging. Images were acquired using a Leica SP8 confocal microscope (Leica microsystems) with appropriate spectral windows for mVenus, images were processed with Fiji software.

### Locomotor behaviour

Adult female and male flies were collected shortly after eclosion and separated into 10 cohorts consisting of 10 flies (100 total) for each genotype. Flies were maintained at 25°C and transferred to fresh food every three days. For the climbing assay, each cohort was transferred to 10ml serological pipette, and allowed to acclimatize for five min. For each trial, flies were tapped down to the bottom of the vial, and the percentage of flies able to cross a five-ml mark successfully within 10 seconds was recorded as the climbing index. Five trials were performed for each cohort, with a 1-min recovery period between each trial. Climbing assays were performed 10 days after eclosion.

### Drosophila larval injections

Injections were performed by using the Nanoliter 2000 (World Precision Instruments, Hertfordshire, United Kingdom) mounted on a micromanipulator (Narishige, London, United Kingdom). Micropipettes were pulled from glass capillary tubes (1.14 mm OD, 0.530 mm ± 25 μm ID; #4878, WPI) using a laser-based micropipette puller (Sutter P-2000, Sutter Instrument, Novato, CA, USA). Third instar larvae (wandering stage) were transferred to an adhesive surface after being quickly washed with water to remove food residues and gently dried using paper tissue. The micropipette was positioned over the approximate centre of the body, on the dorsal side, and the tip was advanced through the cuticle into the hemocoel of the larva. Larvae were injected with 69 nL of PBS (phosphate-buffered saline) supplemented with 10% (v/v) filtered food dye (PME, moss green food colouring; 0.2 µm filter). Food dye was included to aid in monitoring the success of the injection under a dissection microscope (Leica MZ65, Milton Keynes, United Kingdom). ω-hexatoxin-Hv1a (Hv1a, Syngenta, Schaffhauserstrasse, CH-4332 Stein, Switzerland) and α-Bungarotoxin α-Btx (ab120542, Abcam, Cambridge, United Kingdom) were added to the injection mix in order to obtain a final concentration of 2.5 nmol/g and 1.25 nmol/g, respectively (average larval weight was 2.14 mg). After injection, larvae were then gently transferred into agar/grape juice (Ritchie Products Limited, Burton-On-Trent, United Kingdom) plates and kept at 25°C. The rate of survival (expressed as percentage) was calculated as the number of living pupae, formed 1-2 days after injection, divided by the total number of injected larvae. Experiments were repeated three times independently with a total number of 10 larvae for each experimental group. Results were analysed with One-way ANOVA followed by Bonferroni’s multiple comparisons test using GraphPad Prism (version 7, GraphPad Software, San Diego, California, USA).

### Coupling procedure of α-Bungarotoxin to affinity beads

Coupling of α-Bungarotoxin, α-Btx (ab120542, Abcam, Cambridge, United Kingdom) to cyanogen bromide-activated (CNBr) sepharose beads 4B (C9 142-5G, Sigma-Aldrich, Haverhill, United Kingdom) was performed as described (Wang et al., 2003; Mulcahy et al., 2018). CNBr-activated sepharose 4B beads (0.25 g) were hydrated in 1.25 ml of 1 mM HCl for 1 hr at 4°C on a rotator. Beads were centrifuged for 5 min at 1500 × g, the supernatant removed and beads washed twice with 1 ml of coupling buffer (0.25 M NaHCO_3_, 0.5 M NaCl, pH 8.3). Beads were centrifuged for 5 min at 1500 × g and the supernatant was removed. Alpha-Btx (1 mg) was resuspended in 1 ml coupling buffer and incubated together with the affinity beads at 4°C for 16 hr on a rotator. Beads were centrifuged for 5 min at 1500 × g. Coupling efficiency was determined using a Pierce^TM^ quantitative fluorometric peptide kit and used according to the manufacturer’s instructions (23290, Thermo Scientific^TM^, Bishop’s Stortford, United Kingdom). Beads were blocked with 1 ml of 0.2 M glycine in 80 % coupling buffer at 4°C for 16 hr on a rotator. Beads were then centrifuged for 5 min at 1500 × g and washed with 1 ml of 0.1 M NaHCO_3_, 0.5 M NaCl, pH 8.0. This step was repeated with 1 ml of 0.1 M NaCH_3_CO_2_, 0.5 M NaCl, pH 4.0. Beads were washed again in 1 ml of 0.1 M NaHCO_3_, 0.5 M NaCl, pH 8.0. After a final wash step with 1 ml coupling buffer the beads were incubated twice for 30 min in 1 ml Tris-buffer (50 mM Tris, 150 mM NaCl, pH 8.0). The beads were centrifuged for 5 min at 1500 × g, the supernatant was removed and 20 μl Tris-buffer, pH 8.0 was added.

### Membrane protein enrichment and incorporation in SMALPs

*D. melanogaster* heads were obtained and separated according to (Depner et al., 2014). In a 50 ml falcon tube approximately 6 g flies were rapidly frozen in liquid nitrogen and vortexed twice for 3 min, with the tube cooled for 30 sec in liquid nitrogen between. Heads were separated from bodies by sieving (1201124 & 1201125, Endecotts, London, United Kingdom). 1 ml of isotonic lysis buffer (0.25 M sucrose, 50 mM TRIS/HCl pH 7.4, 10 mM HEPES pH 7.4, 2mM EDTA, Protease inhibitor) was added to approximately 0.8 g separated heads. The solution was mixed three times by vortexing and the heads were lysed with 60 strokes in a Dounce homogenizer with a pestle. Membrane protein preparation was performed by differential centrifugation-based fractionation as described (Depner et al., 2014; Geladaki et al., 2019).

Membrane protein pellets were resuspended in 20 to 100 μl 5 % SMALP solution (5 % styrene maleic acid copolymer (3:1), 5 mM Tris-Base, 0.15 mM NaCl, pH 8.0). For efficient incorporation and formation of SMALPs, membrane proteins were incubated with 5 % SMALP solution for 2 hr at room temperature on a rocking platform. To separate the insoluble proteins from the soluble SMALPs a centrifugation step at 100000 × g for 60 min, 4°C was performed. Supernatant containing the SMALPs was combined and used for the nAChRs pull-downs.

### Enrichment of nAChRs by α-Btx pull-down

SMALPs (20-35 mg/ml) were incubated with 200 μl α-Btx conjugated affinity beads for 16 hr, 4°C on a rotator. The beads were then centrifuged for 5 min at 1500 × g and washed two or three times, each for 10 min with 1 ml ice-cold TBS (50 mM Tris, 150 mM NaCl, pH 8.0) on a rotator at 4°C. Beads were centrifuged for 5 min at 1500 × g and nAChRs selectively eluted twice with 100 μl 1 M carbachol (CAS 51-83-2, Insight Biotechnology Ltd, Wembley, United Kingdom). These steps were performed for 25 min at room temperature on a rotator. Beads were centrifuged for 5 min at 1500 × g and eluates were combined and ice-cold 100 % acetone in the volume of four times of the sample was added to the samples, mixed by vortexing and proteins were precipitated for 16 hr at -20°C. Samples were centrifuged at 13000 × g for 15 min. Supernatant was removed and dried proteins were dissolved in Laemmli buffer (1M Tris pH 6.8, 10 % SDS, 5 % glycerol, 2 % bromophenol blue). Proteins were heated at 60°C and loaded on Mini-Protean TGX precast gels (456–1084, 4-15 %, Bio-Rad Laboratories, Inc., Watford, United Kingdom).

### Electron microscopy preparation

For negative staining analysis, membrane proteins were extracted with 5 % SMA and nAChRs were enriched using α-Btx affinity pull-downs. Proteins were diluted 1:10 with deionised water to approximately 0.9 mg/ml and an aliquot of the samples were absorbed onto a glow-discharged copper/carbon-film grid (EM Resolutions) for approximately 2 min at room temperature. Grids were rinsed twice in deionised water and negative staining was performed using a 2 % aqueous uranyl acetate solution. Samples were viewed in a Tecnai G2 transmission electron microscope (TEM, FEI/ThermoFisher) run at 200 keV accelerating voltage using a 20 μm objective aperture to increase contrast; images were captured using an AMT CCD camera.

### Sample preparation for liquid chromatography–mass spectrometry (LC-MS)

The protein lanes were excised from the gels and proteolytic digestion with trypsin/lys-C mix (V5073, Promega, Southampton, United Kingdom) was performed as described (Shevchenko et al., 2007). The gel pieces were covered with 50 mM NH₄HCO₃ / 50 % ACN and shaken for 10 min. This step was repeated with 100 % acetonitrile and finally dried in a speed vac. Samples were reduced with 10 mM DTT in 50 mM NH₄HCO₃ at 56°C for 1 hr and alkylated with 50 mM iodoacetamide in 50mM NH₄HCO₃ at room temperature without light for 45 min. The gels were covered with 50 mM NH₄HCO₃ and 100 % ACN and shaken for 10 min. These steps were repeated and samples were dried in a speed vac. Trypsin/lys-C buffer was added to the sample according to manufacturer’s instructions and incubated for 45 min on ice. Next 30 μl 25 mM NH₄HCO₃ was added and samples were incubated at 37°C for 16 hr. The gel pieces were covered with 20 mM NH₄HCO₃ and shaken for 10 min. Supernatant with peptides was collected. Next, the gels were covered with 50 % ACN / 5 % FA and shaken for 20 min. These steps were repeated and peptides were dried in a speed vac. Samples for glycopeptide enrichment were digested in-solution according to (Queiroz et al., 2019). Samples were reduced and alkylated in 10 mM DTT and 50 mM iodoacetamide. Proteins were digested in final concentration of 2.5 μg trypsin/lys-C buffer for 16 hr at 37°C.

### Peptide clean-up

Peptides were desalted using C-18 stage tips according to (Rappsilber et al., 2007). C-18 material (three C-18 plugs were pasted in a 200 μl pipette tip, Pierce^TM^ C18 Spin Tips, 84850 Thermo Scientific^TM^, Bishop’s Stortford, United Kingdom) was equilibrated with methanol/0.1 % FA , 70 % ACN/0.1 % FA and with 0.1 % FA. Peptides were loaded on C-18 material, washed with 0.1 % FA and eluted with 70 % ACN/0.1 % FA. Samples were dried and finally, peptides were resuspended in 20 μl 0.1 % FA. For glycopeptide enrichment peptides were first desalted using poros oligo r3 resin (1-339-09, Thermo Scientific^TM^, Bishop’s Stortford, United Kingdom) as described (Gobom et al., 1999; Queiroz et al., 2019). Pierce™ centrifuge columns (SH253723, Thermo Scientific^TM^, Bishop’s Stortford, United Kingdom) were filed with 250 μl of poros oligo r3 resin. Columns were washed three times with 0.1 % TFA. Peptides were loaded onto the columns and washed three times with 0.1 % TFA and subsequently eluted with 70 % ACN.

### Glycopeptide enrichment

Enrichment of glycopeptides of nAChRs was performed as described (Hägglund et al., 2004). Micro columns were prepared with 200 μl peptide tips filled with a C8 plug and iHILIC – fusion 5µm, 100 Å silica based material (HCS 160119, Hilicon, Umeå, Sweden). Peptides were solubilized stepwise in 19 μl dH_2_O and then in 80 μl ACN plus 1 μl TFA acid. The micro columns were cleaned with 50 μl 0.1 % TFA and three times equilibrated with 100 μl 80 % ACN, 1 % TFA. Peptides were loaded onto the micro column and washed twice with 100 μl 80 % ACN, 1 % TFA. Glycopeptides were eluted from the column using twice 40 μl 0.1 % TFA and finally with 20 μl 80 % ACN, 1 % TFA. Samples were dried in a speed vac before peptides were deglycosylated with Endo H or PNGase F according to manufacturer’s instructions (P07025 & P0710S, New England Biolabs Inc., Hitchin, United Kingdom).

### LC-MS/MS

Peptide samples were dissolved in 20 μl of 0.1 % (v/v) FA. Approximately 1 μg peptide solution was used for each LC-MS/MS analysis. All LC-MS/MS experiments were performed using a Dionex Ultimate 3000 RSLC nanoUPLC (Thermo Fisher Scientific Inc, Waltham, MA, USA) system and a Q Exactive^TM^ Orbitrap mass spectrometer (Thermo Fisher Scientific Inc, Waltham, MA, USA). Separation of peptides was performed by reverse-phase chromatography at a flow rate of 300 nL/min and a Thermo Scientific reverse-phase nano Easy-spray column (Thermo Scientific PepMap C18, 2μm particle size, 100A pore size, 75 μm i.d. x 50 cm length). Peptides were loaded onto a pre-column (Thermo Scientific PepMap 100 C18, 5μm particle size, 100A pore size, 300 μm i.d. x 5mm length) from the Ultimate 3000 autosampler with 0.1 % FA for 3 min at a flow rate of 15 μL/min. After this period, the column valve was switched to allow elution of peptides from the pre-column onto the analytical column. Solvent A was water + 0.1 % FA and solvent B was 80 % ACN, 20 % water + 0.1 % FA. The linear gradient employed was 2-40 % B in 90 min (the total run time including column washing and re-equilibration was 120 min). In between runs columns were washed at least four times to avoid any carryovers. The LC eluant was sprayed into the mass spectrometer by means of an Easy-spray source (Thermo Fisher Scientific Inc.). An electrospray voltage of 2.1 kV was applied in order to ionize the eluant. All *m/z* values of eluting ions were measured in an Orbitrap mass analyzer, set at a resolution of 35000 and scanned between *m/z* 380-1500 Data dependent scans (Top 20) were employed to automatically isolate and generate fragment ions by higher energy collisional dissociation (HCD, Normalised collision energy (NCE): 25 %) in the HCD collision cell and measurement of the resulting fragment ions were performed in the Orbitrap analyser, set at a resolution of 17500. Singly charged ions and ions with unassigned charge states were excluded from being selected for MS/MS and a dynamic exclusion of 20 seconds was employed.

### Peptide/protein database searching

Protein identification was carried out using sequest HT or mascot search engine software operating in Proteome Discoverer 2.3 (Eng et al., 1994; Koenig et al., 2008). Raw flies were searched against the uniprot *Drosophila_melanogaster*_20180813 database (23297 sequences; 16110808 residues) and a common contaminant sequences database. The search parameters using mascot algorithm were: (i) trypsin was set as the enzyme of choice, (ii) precursor ion mass tolerance 20 ppm, (iii) fragment ion mass tolerance 0.1 Da, (iv) maximum of two missed cleavage sites were set, (v) a minimum peptide length of six amino acids were set, (vi) fixed cysteine static modification by carbamidomethylation, (vii) variable modification by methionine oxidation & deamidation on asparagine and glutamine and N-acetylhexosamine (HexNAc(1)dHex(1) + HexNAc on asparagine) as variable glycopeptide modifications, (viii) A site probability threshold of 75 % was set, (ix) Percolator was used to assess the false discovery rate and peptide filters were set to high confidence (FDR<1).

### Data handling and statistical analysis

Protein data evaluation was performed using R 3.5.3 (Ihaka & Gentleman, 1996). Plotting of graphs were performed in RStudio 1.3.959 (Rstudio, 2020) using ggplot2 (Ginestet, 2011) and other R packages. In order to characterise membrane proteins the following tools were used: (i) TMHMM - 2.0 (Krogh et al., 2001), (ii) PRED-TMBB2 (Tsirigos et al., 2016) (iii) SwissPalm (Blanc et al., 2015), (iv) PredGPI (Pierleoni et al., 2008), (v) Gravy calculator (www.gravy-calculator.de), (vi) Myristoylator (Bologna et al., 2004) (vii) Solubility scores (Sormanni et al., 2015; Sormanni et al., 2017). Analysis of gene ontology (GO) slim terms (The Gene Ontology Consortium 2019) were performed within proteome discoverer 2.3 (Thermo Fisher Scientific). KEGG (Kanehisa et al., 2020) pathway enrichment analysis was performed using DAVID (Huang et al., 2009). For each experimental investigation n ≥ 3 were considered and data are represented as means ± SEM. Experiments were performed in a blinded manner whenever possible. Data are presented as mean ± SD. Statistical tests for SMALPs were performed using two-tailed t-test with an unequal variance and *P* values of ≤ 0.05 were considered to be significant. In DAVID, Fisher’s exact *P* values are computed to measure the gene-enrichment terms. Fisher’s exact *P* value of 0 represents perfect enrichment of a term. Usually *P* value of ≤ 0.05 are to be considered as strongly enriched. In this study the default threshold set in DAVID of 0.1 was used. Linear regression analysis was performed in order to study the efficiency of SMALPs extraction of membrane receptors.

### Structural assessment and illustration of nAChR subunits

For structural alignment of nAChRs matchmaker command operating in UCSF Chimera X 0.91 (Goddard et al., 2018) was used. This command is superimposing protein structures by first creating pairwise sequence alignments, then fitting the aligned residue pairs and displays in an overlaid structure as a result. The following parameters were set to create the aligned structure: (i) alignment algorithm; Needleman-Wunsch (ii) similarity matrix; BLOSUM-62. Structural animation was performed in Blender 2.8 (www.blender.org), an open-source 3D graphics software. For annotation of protein sequences InterProScan was used (Mitchell et al., 2019). Illustrator for biological sequences (IBS) web server was used to present biological sequences (Liu et al., 2015). Multiple sequence alignments were performed (Madeira et al., 2019) or using BoxShade multiple sequence alignments (Swiss institute of bioinformatics).

## Acknowledgements

We thank Professor Tim Dafforn for kindly providing us styrene maleic acid (SMA) copolymer, Dr. Daniel Nightingale for helpful exchange and Mrs Renata Feret for technical discussions. We are very grateful to Syngenta and the Milner Therapeutics Institute for excellent infrastructural support. Electron microscopy was performed using the facilities at CAIC (Cambridge Advanced Imaging Centre, University of Cambridge). Funding was provided by BBSRC (BB/P021107/1) and Syngenta.

## Author Contributions

Conceptualization, BD, DK, CNGG, LC, FGE, SR, and KSL; Methodology, DK, BD, CNGG, RMLQ, GJ, MJD, DPM, and KHM; Data examination, BD, DK, CNGG, LCF, RMLQ, and DPM; Manuscript preparation, BD, DK, CNGG, LCF, FE, SR, and KSL, with contributions of all authors.

## Data availability

The mass spectrometry data from this publication have been deposited to PRIDE (http://www.ebi.ac.uk/pride/archive/) with the data set identifier PXD028484. Biochemical source data is provided (Biochemical_source_data.xls).

## Appendix tables and figures supplement

**Table 1.**
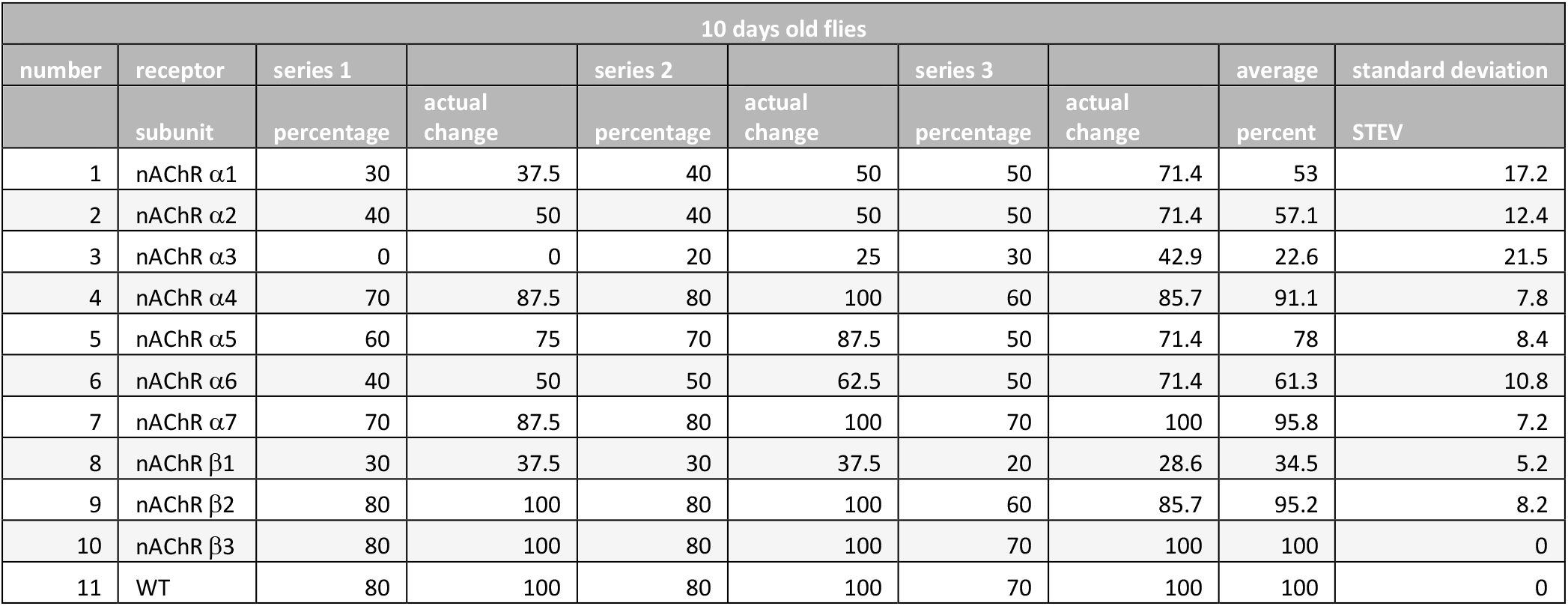
Climbing ability.

| 10 days old flies |  |  |  |  |  |  |  |  |  |
| --- | --- | --- | --- | --- | --- | --- | --- | --- | --- |
| number | receptor | series 1 |  | series 2 |  | series 3 |  | average | standard deviation |
|  | subunit | percentage | actual change | percentage | actual change | percentage | actual change | percent | STEV |
| 1 | nAChR α1 | 30 | 37.5 | 40 | 50 | 50 | 71.4 | 53 | 17.2 |
| 2 | nAChR α2 | 40 | 50 | 40 | 50 | 50 | 71.4 | 57.1 | 12.4 |
| 3 | nAChR α3 | 0 | 0 | 20 | 25 | 30 | 42.9 | 22.6 | 21.5 |
| 4 | nAChR α4 | 70 | 87.5 | 80 | 100 | 60 | 85.7 | 91.1 | 7.8 |
| 5 | nAChR α5 | 60 | 75 | 70 | 87.5 | 50 | 71.4 | 78 | 8.4 |
| 6 | nAChR α6 | 40 | 50 | 50 | 62.5 | 50 | 71.4 | 61.3 | 10.8 |
| 7 | nAChR α7 | 70 | 87.5 | 80 | 100 | 70 | 100 | 95.8 | 7.2 |
| 8 | nAChR β1 | 30 | 37.5 | 30 | 37.5 | 20 | 28.6 | 34.5 | 5.2 |
| 9 | nAChR β2 | 80 | 100 | 80 | 100 | 60 | 85.7 | 95.2 | 8.2 |
| 10 | nAChR β3 | 80 | 100 | 80 | 100 | 70 | 100 | 100 | 0 |
| 11 | WT | 80 | 100 | 80 | 100 | 70 | 100 | 100 | 0 |

**Table 2.** Drosophila larval injection of ω-Hexatoxin-Hv1a & α-Bungarotoxin.

|  | Strain | Injected cpd | Survival (% of pupae formed after injection) |  |  |  |  |
| --- | --- | --- | --- | --- | --- | --- | --- |
|  |  |  | Rep1 | Rep2 | Rep3 | Average | Standard Deviation |
| CONTROL | w<sup>1118</sup> | 2.5 nmol/g Hv1a | 0 | 0 | 0 | 0 | 0 |
| CONTROL - Cas9 lines | THattP40 | 2.5 nmol/g Hv1a | 0 | 0 | 0 | 0 | 0 |
| CONTROL - Cas9 lines | THattP2 | 2.5 nmol/g Hv1a | 0 | 0 | 0 | 0 | 0 |
| nAChR CRISPR mutant | nAChR α1 | 2.5 nmol/g Hv1a | 25 | 0 | 0 | 8.33 | 14.43 |
| nAChR CRISPR mutant | nAChR α2 | 2.5 nmol/g Hv1a | 25 | 0 | 0 | 8.33 | 14.43 |
| nAChR CRISPR mutant | nAChR α3 | 2.5 nmol/g Hv1a | 0 | 0 | 0 | 0 | 0 |
| nAChR CRISPR mutant | nAChR α4 | 2.5 nmol/g Hv1a | 25 | 33.33 | 66.67 | 41.67 | 22.05 |
| nAChR CRISPR mutant | nAChR α5 | 2.5 nmol/g Hv1a | 0 | 0 | 0 | 0 | 0 |
| nAChR CRISPR mutant | nAChR α6 | 2.5 nmol/g Hv1a | 0 | 0 | 0 | 0 | 0 |
| nAChR CRISPR mutant | nAChR α7 | 2.5 nmol/g Hv1a | 0 | 0 | 0 | 0 | 0 |
| nAChR CRISPR mutant | nAChR β2 | 2.5 nmol/g Hv1a | 25 | 33.33 | 66.67 | 41.67 | 22.05 |
| nAChR CRISPR mutant | nAChR β3 | 2.5 nmol/g Hv1a | 0 | 33.33 | 0 | 11.11 | 19.25 |
| Injection CONTROL | w<sup>1118</sup> | PBS | 100 | 100 | 100 | 100 | 0 |
| CONTROL | w<sup>1118</sup> | 1.25 nmol/g α-Btx | 0 | 0 | 0 | 0 | 0 |
| CONTROL - Cas9 lines | THattP40 | 1.25 nmol/g α-Btx | 0 | 0 | 0 | 0 | 0 |
| CONTROL - Cas9 lines | THattP2 | 1.25 nmol/g α-Btx | 0 | 0 | 0 | 0 | 0 |
| nAChR CRISPR mutant | nAChR α1 | 1.25 nmol/g α-Btx | 0 | 0 | 33.33 | 11.11 | 19.25 |
| nAChR CRISPR mutant | nAChR α2 | 1.25 nmol/g α-Btx | 25 | 0 | 33.33 | 19.44 | 17.35 |
| nAChR CRISPR mutant | nAChR α3 | 1.25 nmol/g α-Btx | 0 | 0 | 33.33 | 11.11 | 19.25 |
| nAChR CRISPR mutant | nAChR α4 | 1.25 nmol/g α-Btx | 0 | 33.33 | 0 | 11.11 | 19.25 |
| nAChR CRISPR mutant | nAChR α5 | 1.25 nmol/g α-Btx | 50 | 66.67 | 66.67 | 61.11 | 9.62 |
| nAChR CRISPR mutant | nAChR α6 | 1.25 nmol/g α-Btx | 25 | 66.67 | 66.67 | 52.78 | 24.06 |
| nAChR CRISPR mutant | nAChR α7 | 1.25 nmol/g α-Btx | 50 | 100 | 66.67 | 72.22 | 25.46 |
| nAChR CRISPR mutant | nAChR β2 | 1.25 nmol/g α-Btx | 0 | 33.33 | 0 | 11.11 | 19.25 |
| nAChR CRISPR mutant | nAChR β3 | 1.25 nmol/g α-Btx | 0 | 0 | 0 | 0 | 0 |
| Injection CONTROL | w<sup>1118</sup> | PBS | 100 | 100 | 100 | 100 | 0 |

**Table 3.**
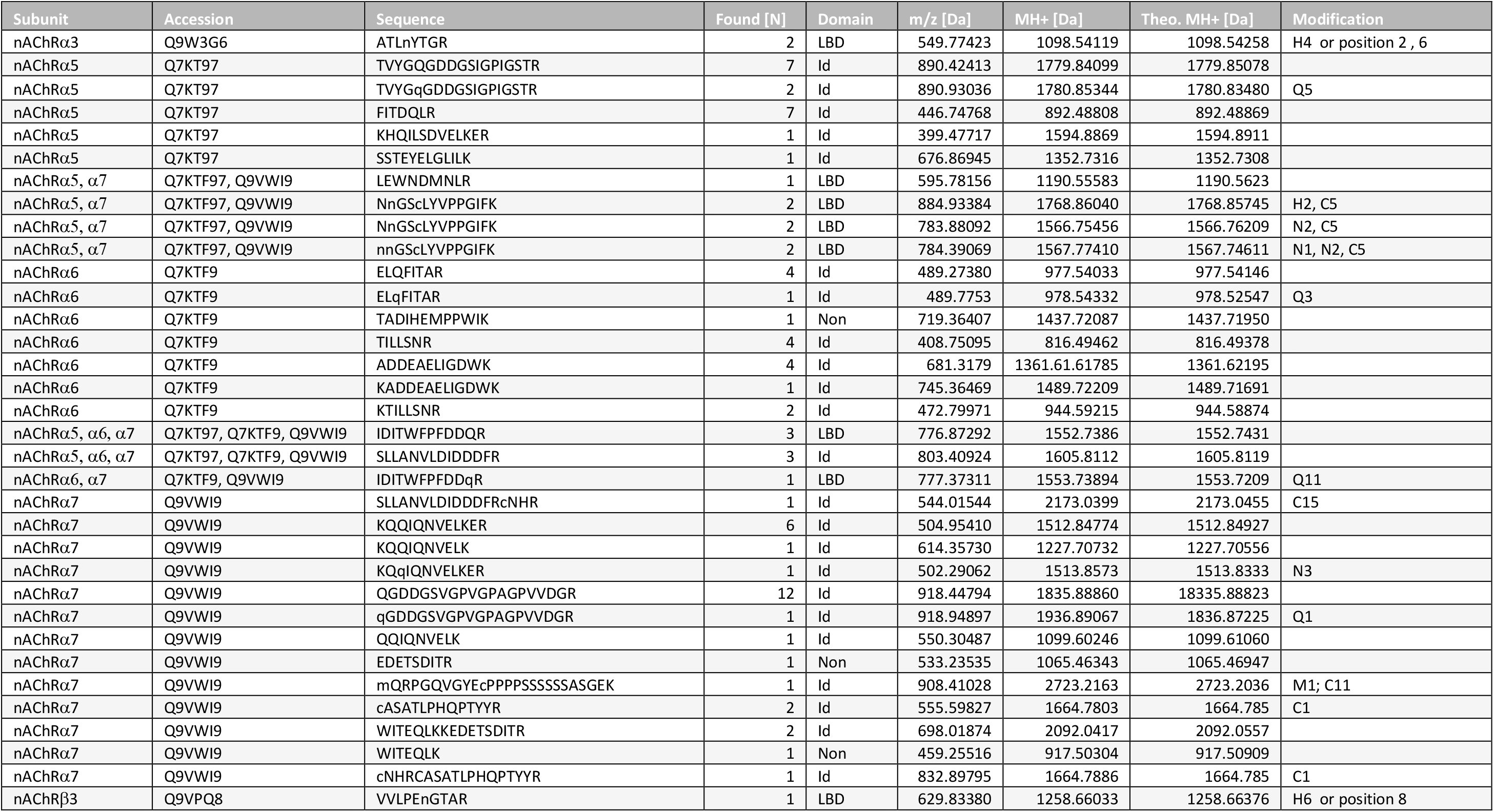
Identified nAChR peptides in pull-downs with α-Bungarotoxin. Peptides from Dα3, Dα5, Dα6, Dα7 and Dβ3 nAChR subunits are listed and found [N] times within individual replicates. Protein domains are marked with: Ed extracellular-, Id Intracellular-, LBD ligand*-* binding-, and Non-domain localization. The mass-to-charge ratio (m/z) of the precursor ions, the protonated monoisotopic masses, the theoretical MH^+^ masses in Dalton [Da] and peptide modifications are listed. Peptide modifications are listed with: (C) Carbamidomethylation; (N,Q) Deamidation; (H) N-acetylhexosamine (HexNAc); (M) Oxidation.

**Table 4.**
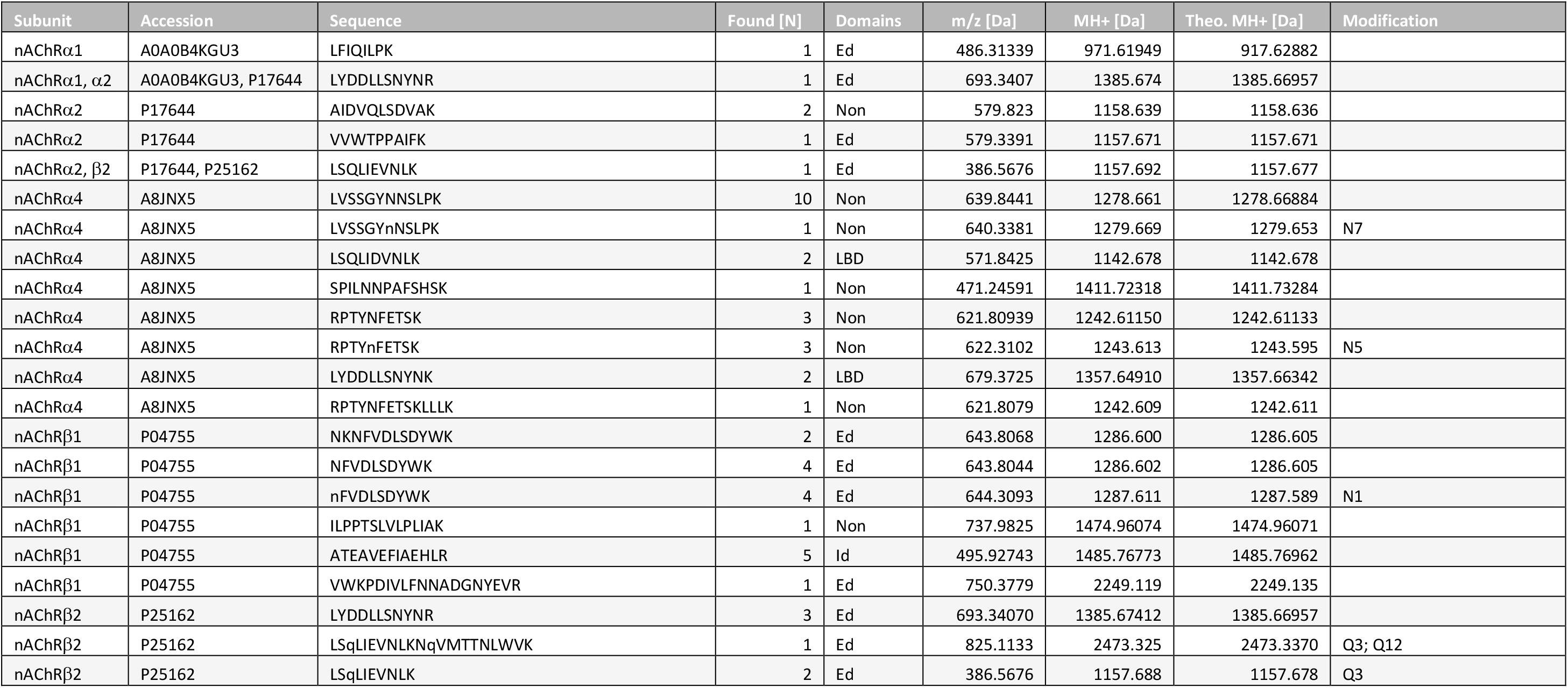
Identified nAChR peptides in pull-downs without α-Bungarotoxin. Identified peptides of nAChR subunits which are found in control pull-down samples without α- Bungarotoxin (α-Btx).

**Table 5.**
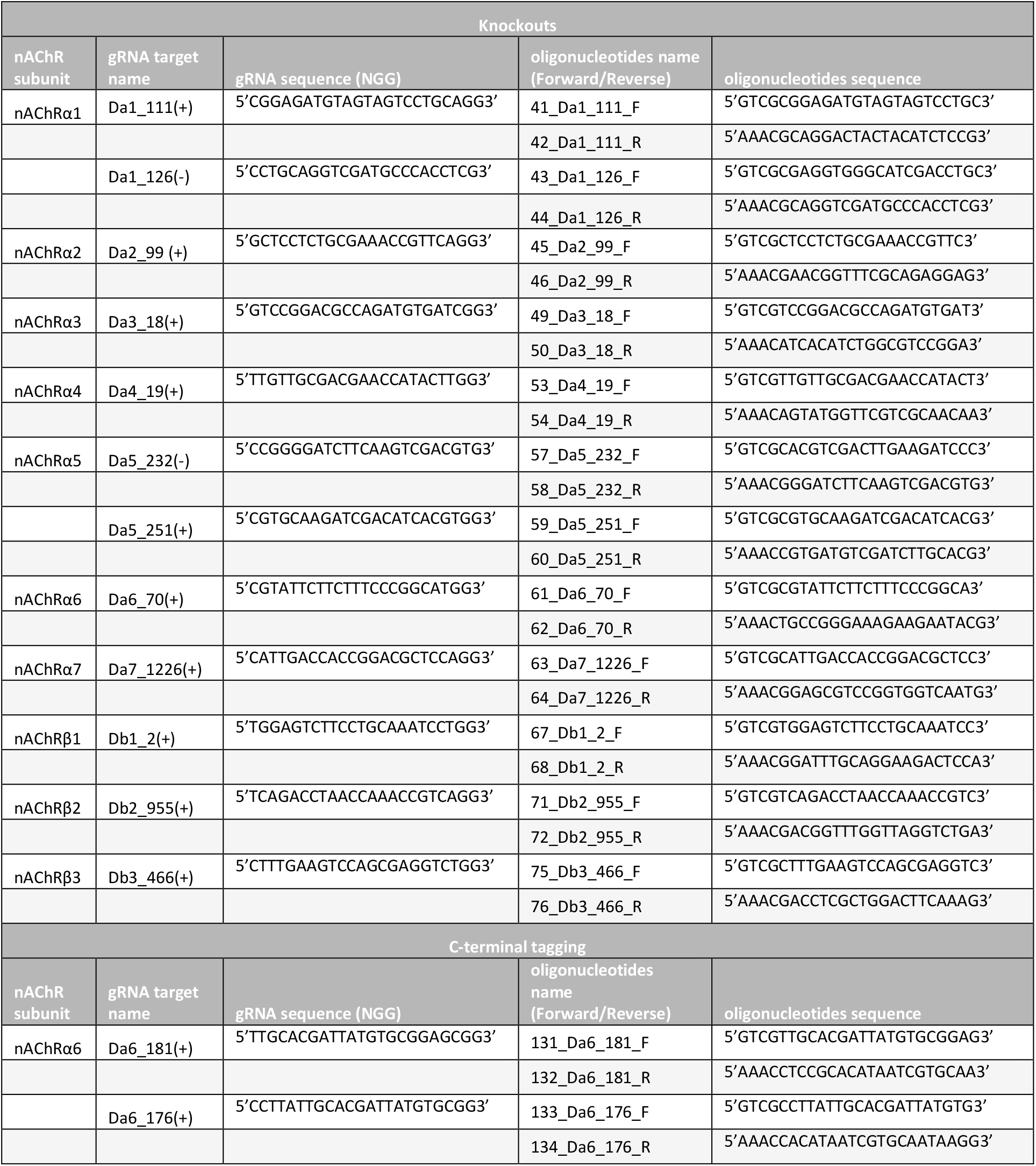
List of gRNAs and oligonucleotides used for cloning.

**Table 6.**
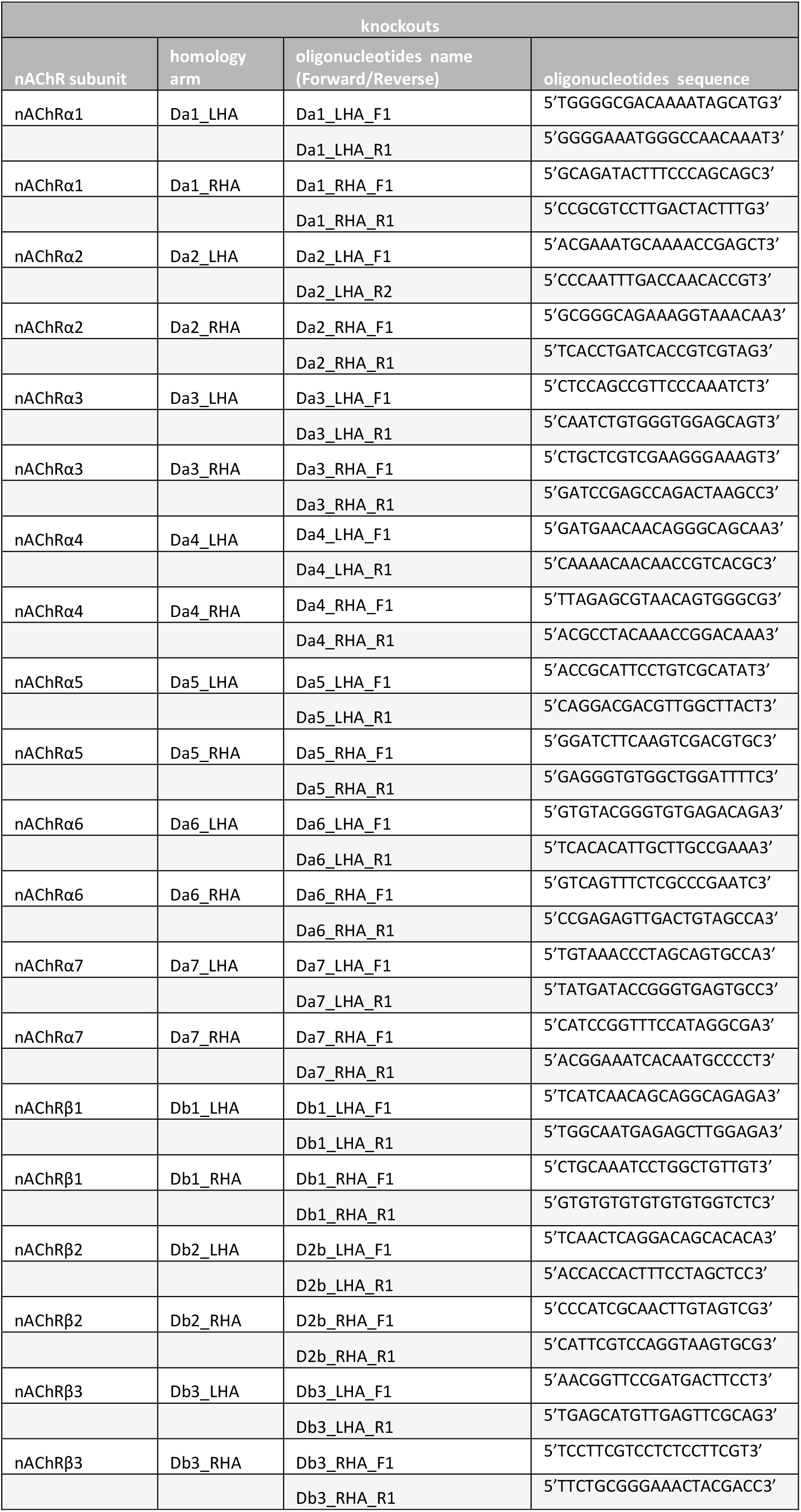
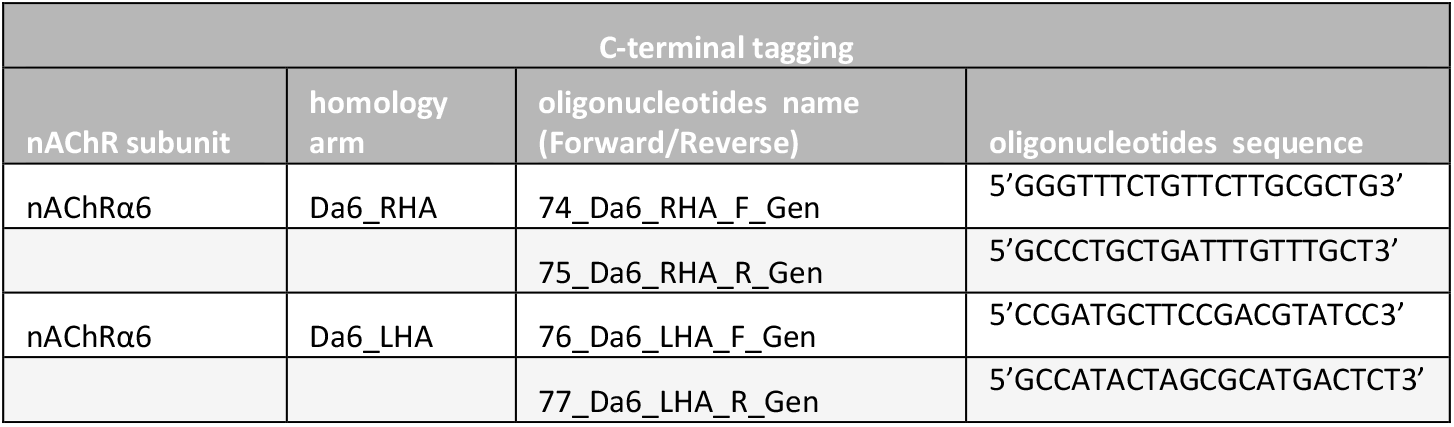
List of oligonucleotides used for amplification from genomic DNA.

**Table 7.**
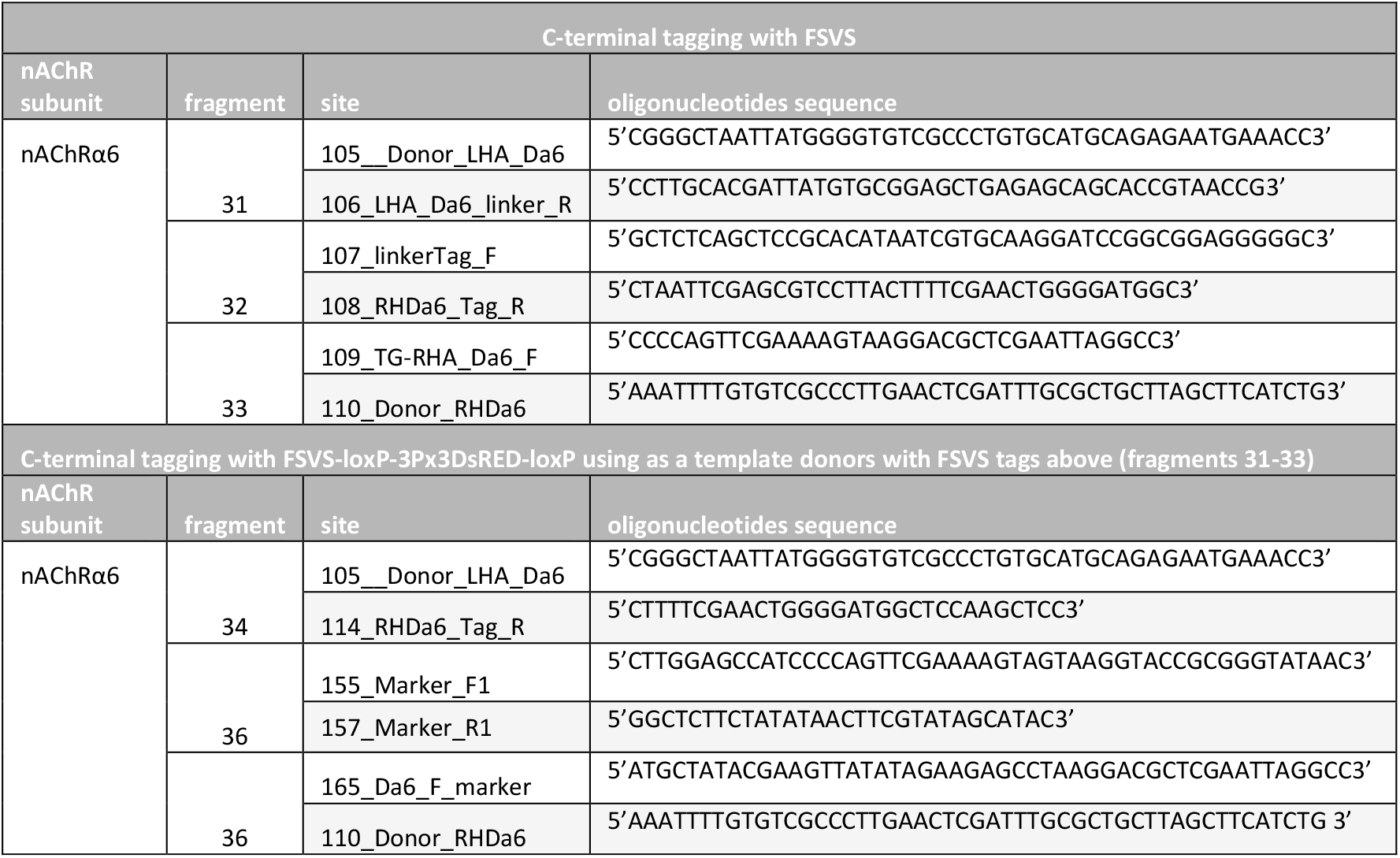
C-terminal tagging of nAChRa6 with FSVS.

**Figure supplement 1.**
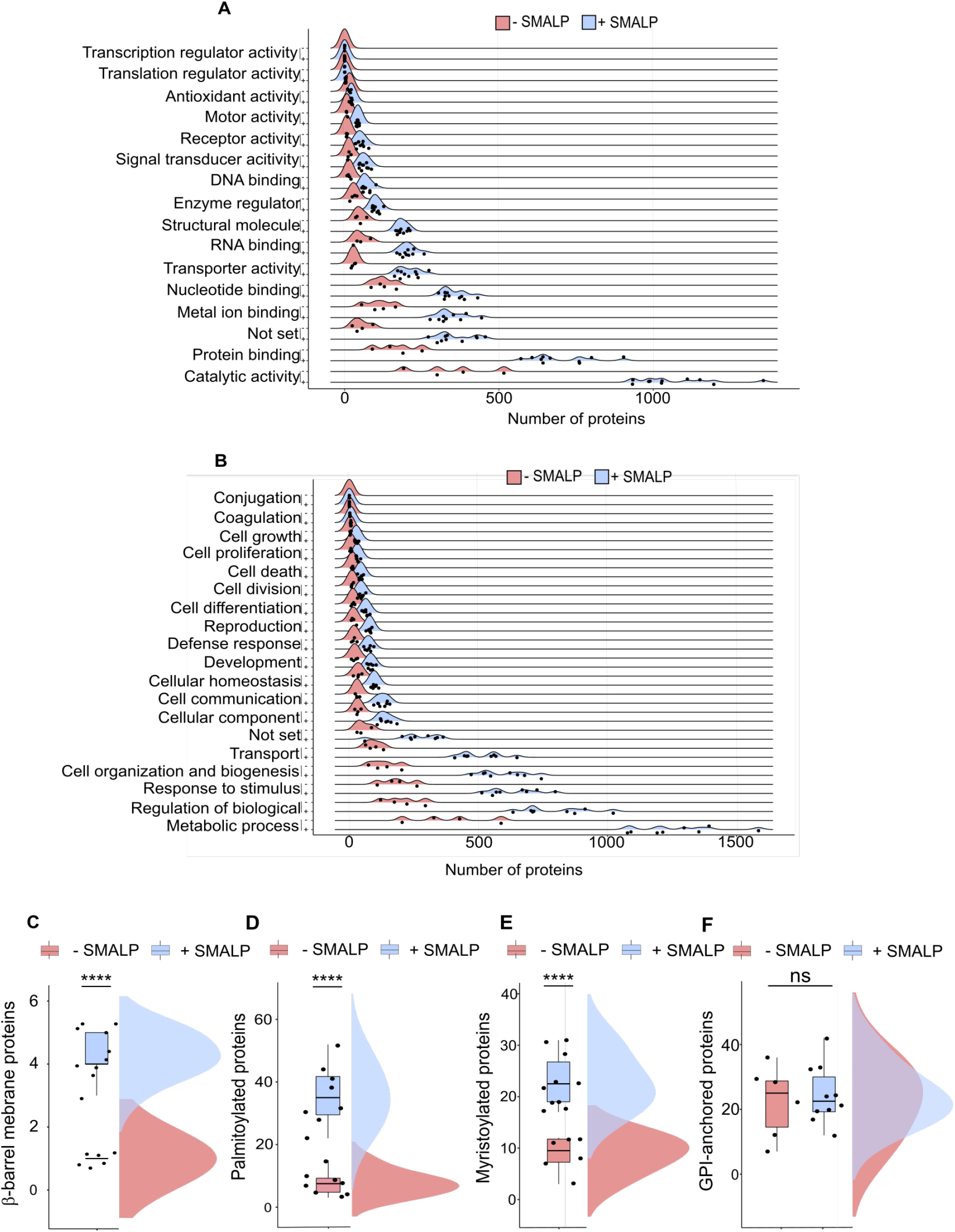
GO terms and predicted membrane proteins. **(A)** GO slim term for biological process and (**B**) for molecular function analysed within samples solubilized without or with SMA, n=4 or 11 per conditions. Predicted β-barrel membrane- (**C**), two-tailed t-test \*\*\*\**P*<0.0001, n=6 or 10; palmitoylated- (**D**), \*\*\*\**P*<0.0001, each n=8; myristoylated- (**E**), \*\*\*\**P*<0.0001, n=6 or 10; and GPI-anchored proteins (**F**), non-significant ns, n=6 or 10.

**Figure supplement 2.**
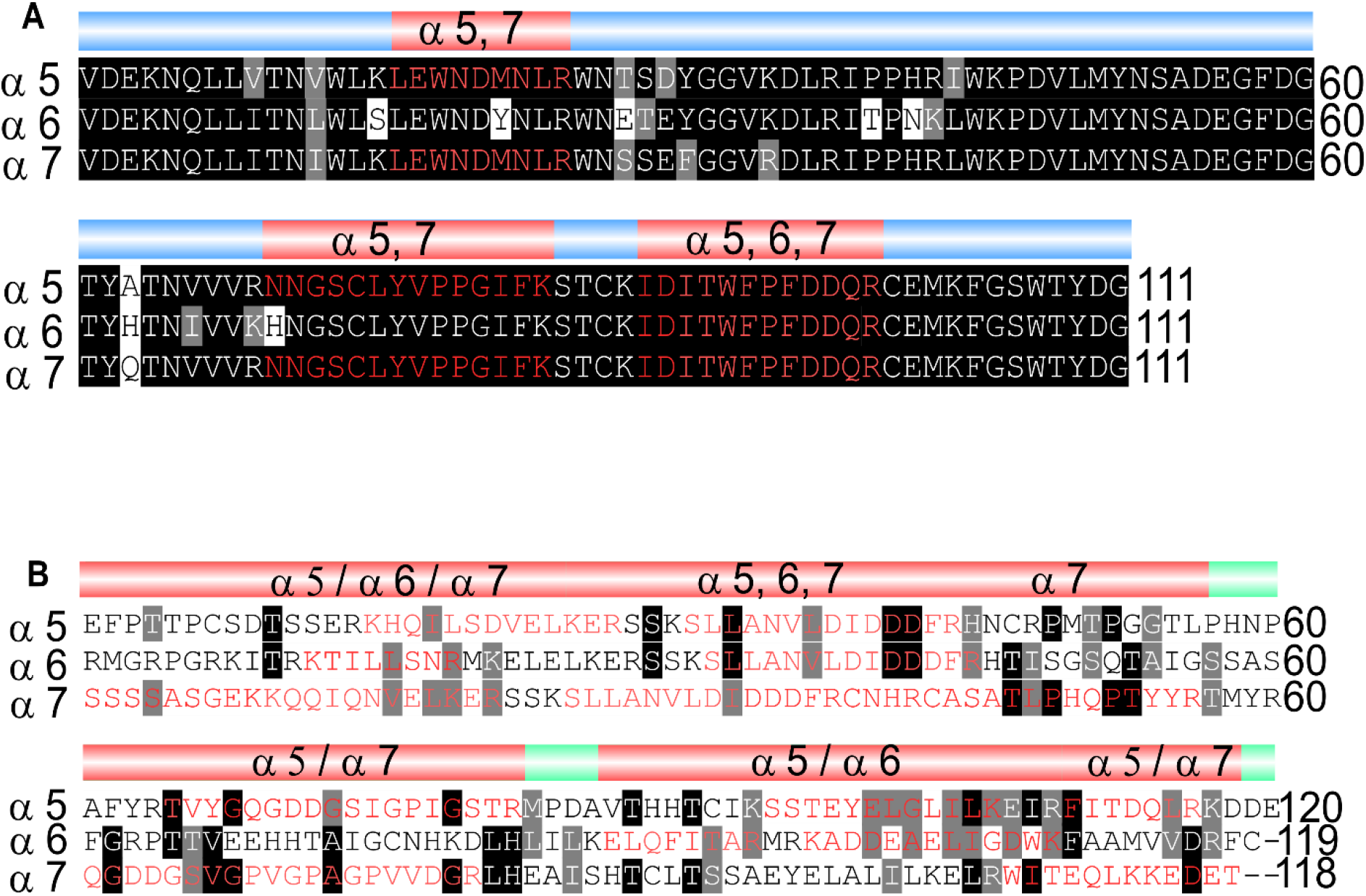
Identified peptides in ligand-binding and cytoplasmic domain. **(A)** Shared peptides found in the ligand-binding domains are shown in red. (**B)** Identified unique (/) and shared (,) peptides in cytoplasmic domains.

**Figure supplement 3.**
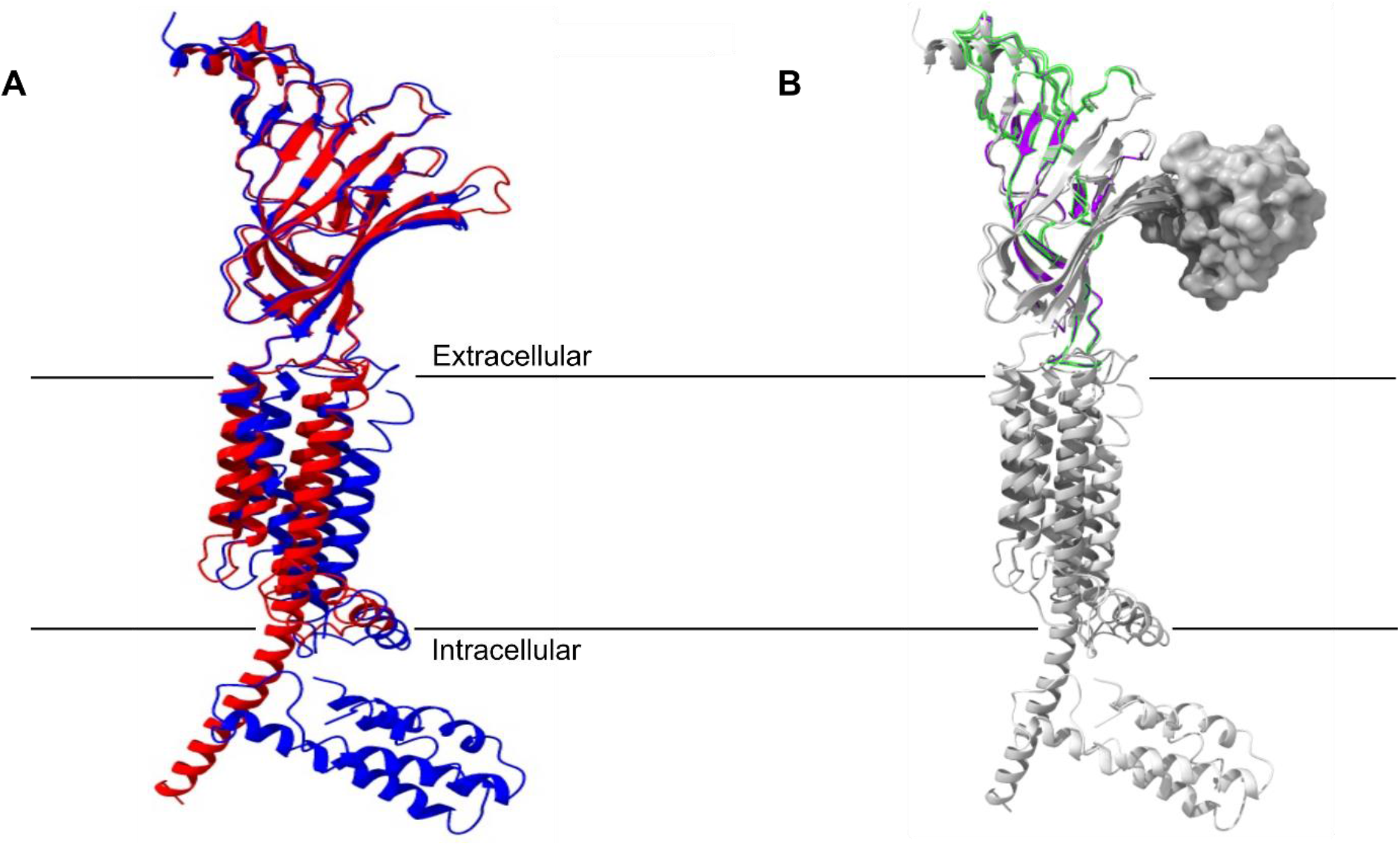
Superimposed nAChR α-subunits structure together with identified peptides. **(A)** Superimposed nAChR α-subunit structures from *Homo sapiens* (blue, PDB 6USF) and *Torpedo californica* (red, 6UWZ). Extracellular ligand*-*binding domain (LBD) illustrates a structure similarity. **(B)** Same superimposed structures bound to α-bungarotoxin (α-Btx, surface structure). Peptides found in LBD are highlighted in green. The homology regions of Dα6 nAChRs LBD are shown in violet.

**Figure supplement 4.**
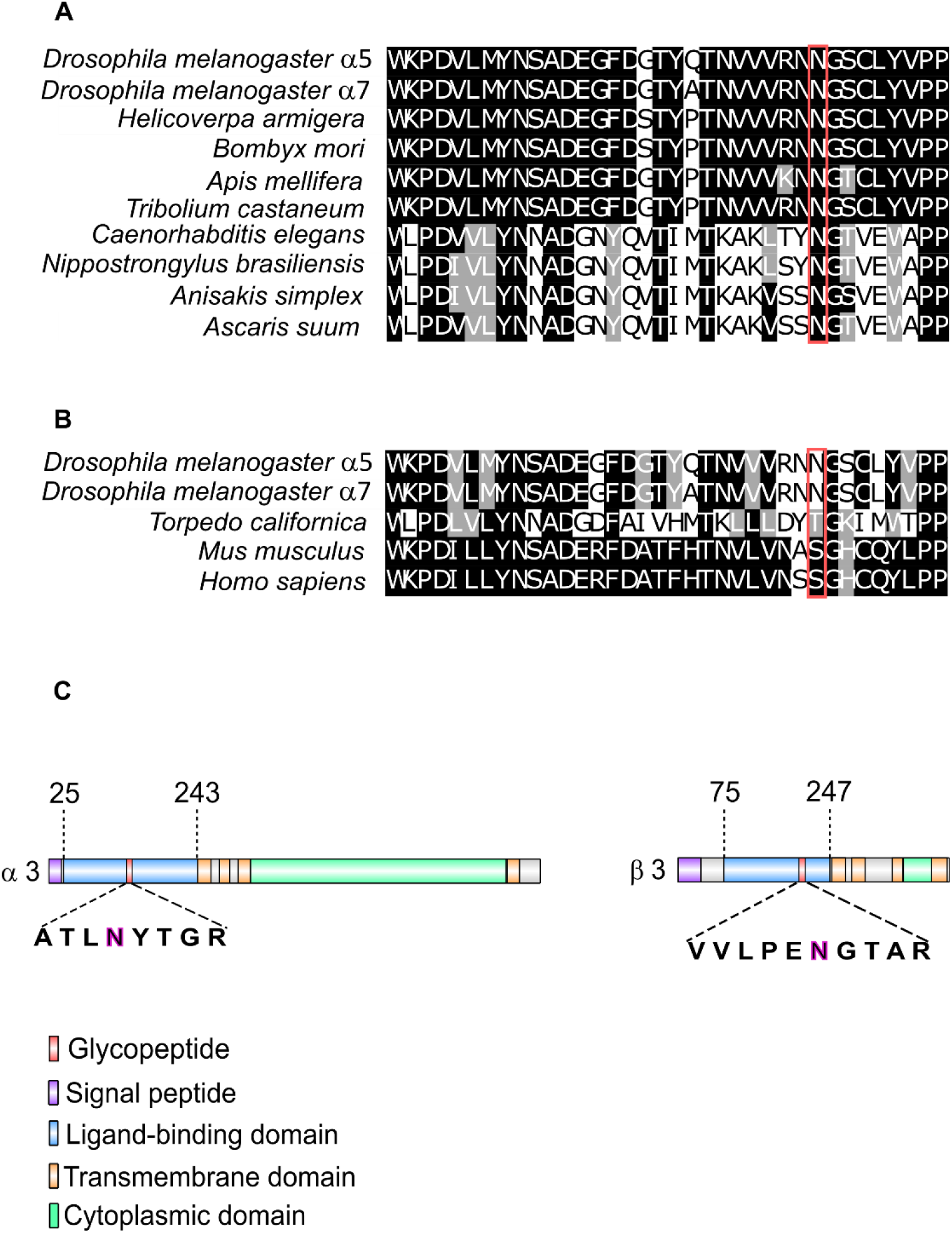
Glycosylation sites of nAChR subunits. **(A)** Multiple sequence alignment of insect α7 nAChR subunits compared to sequences of nematodes. The glycosylated ligand-binding domain (LBD) sequence of Dα5 and Dα7 nAChR subunits are shown. Glycosylated asparagine residues highlighted in red are conserved within insects and nematodes (Dα5 422 and Dα7 170 amino acids). **(B)** Same Dα5 and Dα7 nAChR subunit sequences compared to *T.californica*, *D.rerio*, *M. musculus* and *H. sapiens*. **(C)** Graphical representation of Dα3 and Dβ3 nAChR subunits. N-acetylhexosamine (H) modification on asparagine residues are highlighted and are of low site probability ≤ 80 %.

